# Engineering plant tandem kinase immune receptors expands effector recognition profiles

**DOI:** 10.64898/2025.12.15.694194

**Authors:** Daniel S. Yu, Rafał Zdrzałek, Emi Katayama, Hitomi Akiyama, Lucy Daykin, Nathan J. Williams, Iwan Goodridge, Soichiro Asuke, Mark J. Banfield

## Abstract

Plant intracellular immune receptors are widely deployed in breeding to protect crops from disease. In addition to nucleotide-binding leucine-rich repeat receptors (NLRs), tandem kinase proteins (TKPs) have recently emerged as an important family of immune receptors within staple cereal food crops, but how TKPs recognize effectors and whether they are amenable to engineering is essentially unknown. Here, we show that the barley and wheat TKPs Rmo2 and Rwt7 recognize different blast fungus effectors via their integrated HMA domains using different protein interfaces with nanomolar binding affinity. Structural analysis pinpointed interface residues that dictate effector recognition and enabled engineering of dual-specificity TKPs. These results establish integrated HMA domains as programmable modules within TKPs for designing new specificities in plant immunity for diseases relevant to global agriculture.

## Main Text

Globally, 20–30% of cereal crop production (including wheat, rice and barley) is lost each year to plant diseases (*1-3*). Plant disease defenses are triggered by cell surface or intracellular receptors that recognize pathogen-associated molecular patterns (PAMPs) or effectors released from pathogens during infection (*4*). Structural characterization of intracellular nucleotide-binding leucine-rich repeat receptors (NLRs) (*5*) has elucidated the activation mechanisms for these plant immune receptors, highlighting previously unrecognized potential for engineering disease resistance (*6-10*).

Despite the importance of NLRs, up to 35% of race‐specific resistance genes from wheat and barley do not encode canonical NLRs (*11*). Genes encoding kinase-fusion proteins (KFPs) have emerged as a major source of resistance within cereal crops for multiple agriculturally relevant diseases (*12-24*). Most cloned KFPs are tandem kinase proteins (TKPs), with some of the best characterized examples being Rpg1, Sr62 and RWT4 (*12, 25-27*). While Sr62 and RWT4 are both TKPs, they recognize effectors via different mechanisms. The first kinase domain of Sr62 has a variable beta-finger domain predicted to dimerize and facilitate the binding of AvrSr62 (*25*). RWT4 is predicted to bind AvrPWT4 via its N-terminal kinase duplication (KDup) domain and pseudokinase domain, and substitutions within the KDup domain that prevent effector binding result in disease susceptibility (*27*).

Plant receptors have evolved non-canonical integrated domains that function as effector decoys. For example, a subset of NLRs, such as Pik-1 and RGA5 in rice, have an integrated heavy metal-associated (HMA) domain that mimics the effector host target that facilitates effector binding and immune activation (*28, 29*). Remarkably, integrated HMA domains are also found in TKPs. Rpg1 has an HMA domain at its N-terminus that went unnoticed until recent advancements in structural modeling (*30*). Furthermore, a strikingly similar domain structure was discovered in the TKPs Rmo2 (of barley) and Rwt7 (of wheat), which recognize the effectors PBY2 and PWT7, respectively, from different pathotypes of the blast fungus *Magnaporthe oryzae* (*24*). Whether these domains function to recognize effectors and promote immune responses is unknown.

In this study, we investigated the role of the integrated HMA domains of the TKPs Rmo2 and Rwt7 in effector recognition and initiation of immune signaling. We elucidated the structural and biochemical basis of effector binding and immune signaling specificity. Furthermore, based on the structures, we engineered Rmo2 and Rwt7 with expanded responses that include previously unrecognized effectors. Our work establishes TKP receptors as tools for the generation of new disease-resistance profiles in cereal crops and demonstrates that TKP integrated domain engineering is a viable approach to develop innovative strategies for pathogen control.

### TKP integrated HMA domains directly interact with pathogen effectors *in vitro*

Integrated HMA domains in plant NLRs recognize specific effectors, thereby initiating immune responses (*28, 29*). Similarly, the N-terminal integrated HMA domains of cereal TKP receptors Rmo2 and Rwt7 recognize the *M. oryzae* effectors PBY2 and PWT7, respectively (*24*). The TKP receptors Rmo2, Rwt7, and Rpg1 are highly variable in sequence across their HMA domains, while their pseudokinase and kinase domain sequences are more conserved (fig. S1). Therefore, we hypothesized that the integrated HMA domains of Rmo2 and Rwt7 function as bait effectors, similar to those found in NLR receptors. To test this, we determined the binding specificities and affinities of the integrated HMA domains of Rmo2 and Rwt7 towards PBY2 and PWT7 *in vitro*. We purified recombinant HMA domains (Rmo2^HMA^ and Rwt7^HMA^) and effectors (PBY2 and PWT7) and performed analytical size exclusion chromatography (aSEC). We observed shifts in peak elution time when Rmo2^HMA^ and PBY2, or Rwt7^HMA^ and PWT7 were pre-incubated together, relative to the elution times of the individual proteins, indicating direct interaction. We detected no peak shift when the respective HMA domains were mixed with their non-cognate effectors (fig. S2). Quantitative analysis of the binding affinities using isothermal titration calorimetry (ITC) also demonstrated the specificity of the interaction between Rmo2^HMA^ and PBY2, and between Rwt7^HMA^ and PWT7 (Fig. 1, A and B). These interactions displayed nanomolar binding affinities, with *K*_*D*_ values of 53.5 nM for Rmo2^HMA^ with PBY2 and 2.92 nM for Rwt7^HMA^ with PWT7. Consistent with the aSEC experiments, there was no binding observed between each HMA domain and the non-cognate effector in ITC experiments.

**Fig. 1.**
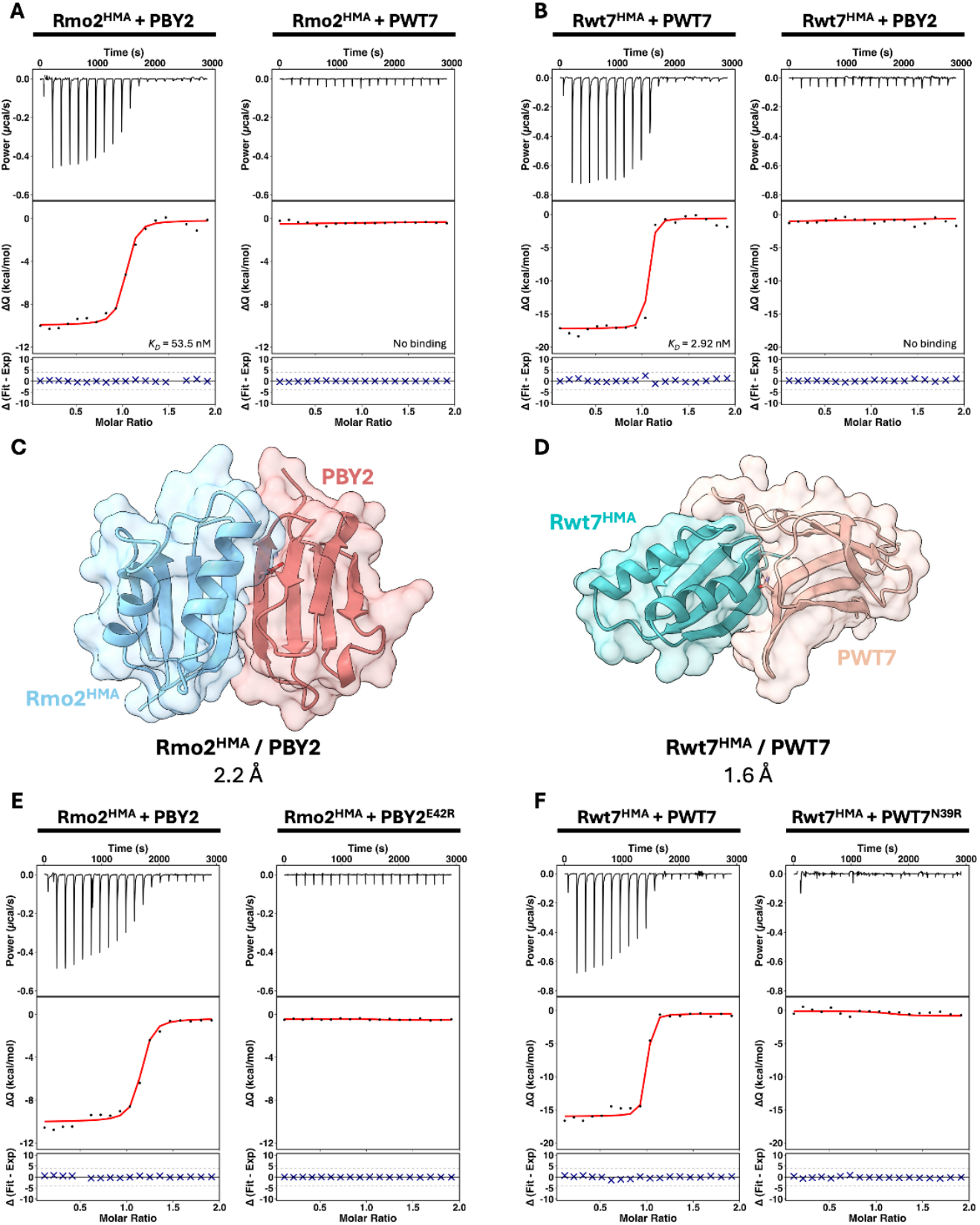
Biophysical and structural analysis of binding specificities from the integrated HMA domains of TKPs. (**A**) Isothermal titration calorimetry (ITC) assay of Rmo2^HMA^ mixed with PBY2 or PWT7, and of (**B**) Rwt7^HMA^ mixed with PWT7 or PBY2. **(C)** Crystal structures of the Rmo2^HMA^/PBY2 (PDB: 9TFO) and (**D**) Rwt7^HMA^/PWT7 (PDB: 9TFP) complexes. Key residues within the effectors involved in interacting with the HMA domains (E42 from PBY2 and N39 from PWT7) are shown as sticks. (**E**) ITC assay of Rmo2^HMA^ mixed with PBY2 or PBY2^E42R^, and of (**F**) Rwt7^HMA^ mixed with PWT7 or PWT7^N39R^. All ITC experiments were performed in triplicate and were analyzed using AFFINImeter.

Barley lines harboring *Rmo2* are resistant to *M. oryzae* pathotypes expressing *PBY2* but not *PWT7*, while wheat lines carrying *Rwt7* are resistant to *M. oryzae* pathotypes expressing *PWT7* but not *PBY2* (*24*). Furthermore, barley protoplasts expressing *Rmo2* and *PBY2*, or *Rwt7* and *PWT7*, exhibit cell-death responses indicative of immune activation, while there is no cell death when receptors and effectors are mismatched (*24*). Replacing Rmo2^HMA^ with Rwt7^HMA^ within Rmo2 causes a switch in recognition specificity from PBY2 to PWT7 in transgenic barley lines (*24*). Collectively, these data demonstrate that the HMA domains of Rmo2 and Rwt7 directly bind their cognate effectors with high affinity and dictate recognition specificity for these receptors.

### Structure-informed interface mutagenesis disrupts protein interactions *in vitro*

To understand the structural basis of the interactions of Rmo2^HMA^ with PBY2 and of Rwt7^HMA^ with PWT7, we determined the structures of their complexes with *X*-ray crystallography. Accordingly, we co-purified and obtained crystals for Rmo2^HMA^/PBY2 and Rwt7^HMA^/PWT7 HMA domain/effector pairs (fig. S3). We solved the crystal structures of Rmo2^HMA^/PBY2 and Rwt7^HMA^/PWT7 by molecular replacement using AlphaFold2 models as templates (*31, 32*), reaching refined resolutions of 2.2 Å and 1.6 Å, respectively (Fig. 1, C and D, table S1). The effectors bound to spatially distinct interfaces on the HMA domains. The interface between Rmo2^HMA^ and PBY2 was predominantly mediated by polar main-chain interactions between the β3 of Rmo2^HMA^ and β2 of PBY2 with only limited polar side-chain interactions (D15 and K18 from Rmo2^HMA^ and E42 and S48 from PBY2) (fig. S4A). By contrast, the Rwt7^HMA^/PWT7 interface displayed many polar side-chain interactions across two faces of Rwt7^HMA^. Central to the interaction interface were the sidechains of N39 and D40 from PWT7, while the first residue of Rwt7^HMA^ (M1) formed extensive interactions with the effector (fig. S4, B and C). We tested whether the interaction interfaces observed in the structures underpin protein–protein interactions by introducing structure-informed single amino-acid substitutions in PBY2 (E42R) and PWT7 (N39R) to disrupt interactions with their respective HMA domains. We tested the binding of each recombinant effector mutant to its cognate HMA domain by aSEC and ITC. No peak shift was observed by aSEC when PBY2^E42R^ and PWT7^N39R^ were incubated with Rmo2^HMA^ or Rwt7^HMA^, respectively (fig. S5). Similarly, each effector mutant showed no quantitative binding to its respective HMA domain by ITC (Fig. 1, E and F). Therefore, structure-informed mutagenesis of the effectors successfully disrupted the interactions with the respective HMA domains *in vitro*.

### Rmo2 and Rwt7 integrated HMA domains are baits for effectors in planta

To establish whether the *in vitro* interactions between HMA domains and effectors reflect effector binding to full-length receptors in plant cells, we used the split GAL4 RUBY assay (*33*). Areas of *Nicotiana benthamiana* leaves co-expressing *Rmo2-GAL4* (encoding Rmo2 fused with the DNA-binding domain of yeast GAL4) and *PBY2-VP16* (encoding PBY2 fused with VP16) accumulated red pigments due to transcriptional activation of betalain production, indicative of protein–protein interactions in this assay (Fig. 2A). Co-expression of *Rmo2-GAL4* and *PWT7-VP16* did not lead to significant betalain accumulation, confirming the specificity of the effector interactions. Furthermore, leaves co-expressing *Rmo2-GAL4* with the effector mutant *PBY2*^*E42R*^*-VP16* did not accumulate betalain, showing that the E42R mutation in PBY2 disrupted interaction with Rmo2 in plant cells. Accumulation of proteins was confirmed in plant tissues by immunoblot (fig. S7). We attempted the split GAL4 RUBY assay with Rwt7-GAL4, but detected little betalain production for any combination, suggesting that Rwt7 may not be suitable for this assay (fig. S6). As replacing the HMA domain of Rmo2 with Rwt7^HMA^ switches recognition specificity of Rmo2 from PBY2 to PWT7 (*24*), we generated the *Rmo2*^*Rwt7-HMA*^*-GAL4* construct and co-expressed it with *PWT7-VP16, PBY2-VP16*, or *PWT7*^*N39R*^*-VP16* (Fig. 2B). We observed significantly higher betalain production in the presence of PWT7 than with PBY2 or PWT7^N39R^, suggesting that the chimeric receptor interacts with PWT7 instead of PBY2, and that the N39R mutation in PWT7 disrupts interaction with Rmo2^Rwt7-HMA^.

**Fig. 2.**
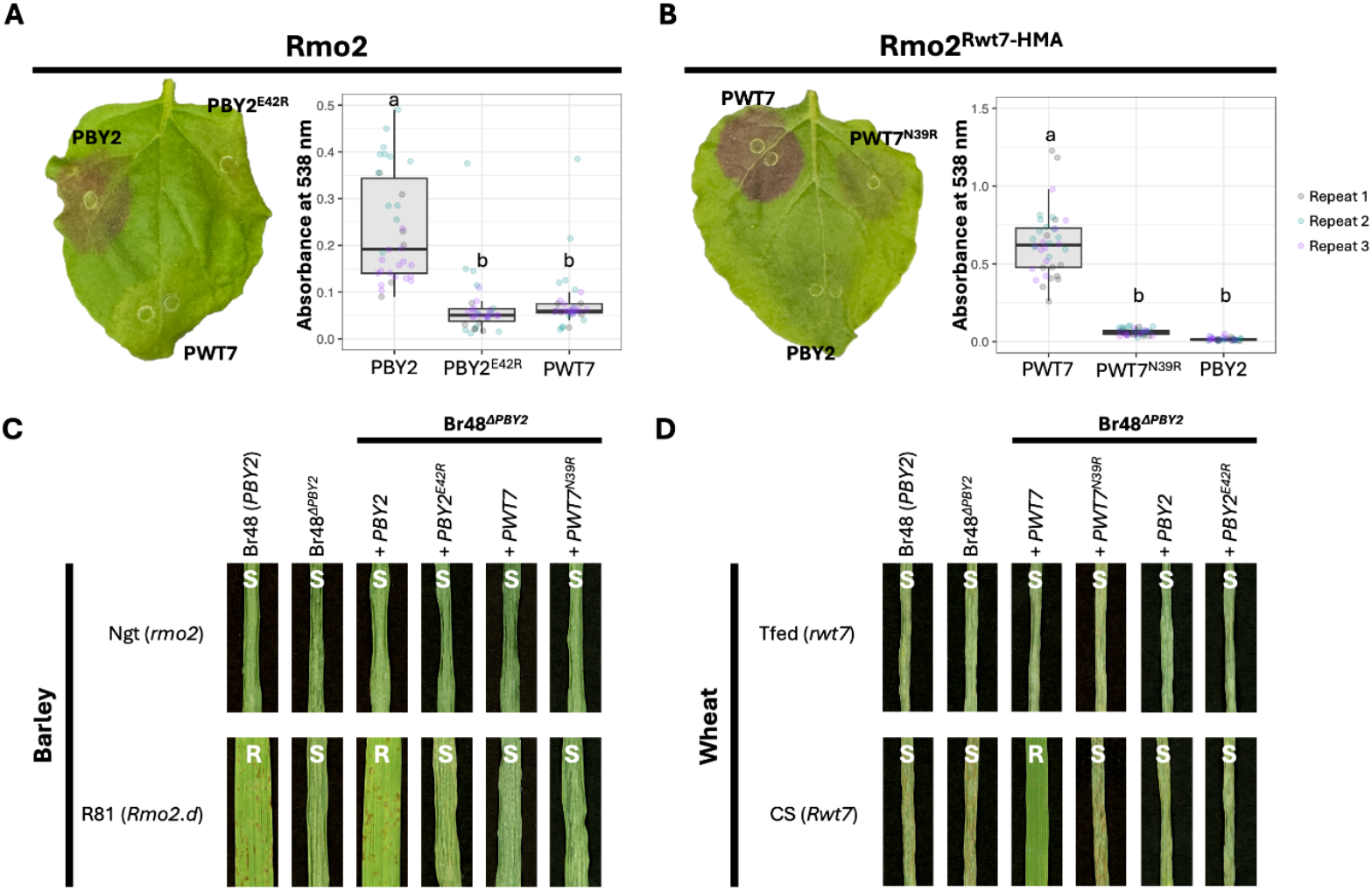
Structure-informed effector mutants no longer bind to or recognize their cognate TKP *in planta*. (**A, B**) Split GAL4 RUBY assay showing betalain production in *Nicotiana benthamiana* leaves following co-expression of **(A)** *Rmo2-GAL4* (encoding Rmo2 with the DNA-binding domain of yeast GAL4 in its C-terminus) with *PBY2-VP16, PBY2*^*E42R*^*-VP16*, or *PWT7-VP16* (encoding the effectors fused to the activation domain of VP16 at their C termini) or (**B**) *Rmo2*^*Rwt7-HMA*^*-GAL4* with *PWT7-VP16, PWT7*^*N39R*^*-VP16*, or *PBY2-VP16*. All expression cassettes were under the control of the cauliflower mosaic virus (CaMV) 35S promoter. Leaves were harvested 4 days after infiltration and chlorophyll was cleared with ethanol. Betalain was extracted from each infiltrated spot and absorbance was measured at 538 nm. Each experiment consisted of 6 or 12 replicates, performed three times independently. Different letters indicate significant differences, as determined by one-way ANOVA and *post-hoc* Tukey’s honestly significant difference tests at *P <* 0.05. (**C, D**) Infection assay with *Magnaporthe oryzae* BR48 derived and complementation isolates on the barley cultivars ‘Nigrate’ (Ngt) and ‘Russia 81’ (R81), or (**D**) on the wheat cultivars ‘Transfed’ (Tfed) and ‘Chinese Spring’ (CS). Primary leaves of eight-day-old seedlings were spray-inoculated with *M. oryzae* isolates. Leaves were imaged 4 days post inoculation. Resistant (R) or susceptible (S) phenotypes are labeled.

### The integrated HMA domains of Rmo2 and Rwt7 bind effectors for immunity in cereals

To link direct receptor–effector interactions to susceptibility or resistance phenotypes during infection, we performed spray inoculation assays on barley and wheat cultivars, challenging them with *M. oryzae* strains expressing effectors or effector mutants (Fig. 2, C and D). The barley cultivar ‘Nigrate’ (Ngt, lacking *Rmo2*) was susceptible to *M. oryzae* strain Br48, which harbors *PBY2*, but the barley cultivar ‘Russia 81’ (R81, carrying *Rmo2*) was resistant (Fig. 2C). The *PBY2* deletion strain (Br48^ΔPBY2^) causes disease on R81 (*24*). Indeed, R81 challenged with Br48^ΔPBY2^ displayed susceptibility, as did R81 challenged with Br48^ΔPBY2^ complemented with *PBY2*^*E42R*^ (Fig. 2C). R81 was resistant to Br48^ΔPBY2^ complemented with *PBY2*. The Ngt and R81 cultivars were both susceptible to Br48^ΔPBY2^ complemented with *PWT7* or *PWT7*^*N39R*^. The wheat cultivars ‘Transfed’ (Tfed, lacking *Rwt7*) and ‘Chinese Spring’ (CS, containing *Rwt7*) were both susceptible to infection with Br48^ΔPBY2^ (lacking *PWT7*) (Fig. 2D). CS was resistant to Br48^ΔPBY2^ complemented with *PWT7*, but susceptible to Br48^ΔPBY2^ complemented with *PWT7*^*N39R*^. Tfed and CS were both susceptible to Br48^ΔPBY2^ complemented with *PBY2* or *PBY2*^*E42R*^.

We conclude that disrupting the interaction between cognate effectors and their corresponding receptor HMA domains prevents recognition by Rmo2 and Rwt7, confirming the critical role of the HMA domain as an integrated effector bait domain within these receptors.

### Structure-informed resurfacing of TKP integrated HMA domains expands their binding profiles

Rmo2^HMA^ and Rwt7^HMA^ specifically interact with PBY2 and PWT7, respectively (Fig. 1, A and B). In the crystal structures, the HMA domains of Rmo2 and Rwt7 interacted with their cognate effectors via unique, spatially distinct interfaces (fig. S8). We thus wondered if resurfacing of the Rmo2 and Rwt7 integrated HMA domains could switch effector specificity and generate a receptor that could bind and respond to both PBY2 and PWT7.

We resurfaced Rmo2^HMA^ along the Rwt7^HMA^/PWT7 interface, changing residues in Rmo2^HMA^ to those found in Rwt7^HMA^, generating Rmo2^HMA+^ (Fig. 3A). ITC assays confirmed that engineered Rmo2^HMA+^ retains nanomolar binding affinity to PBY2, as did native Rmo2^HMA^ (Fig. 3B). Then we tested whether Rmo2^HMA+^ could bind to PWT7. In contrast to Rmo2^HMA^, which showed no binding to PWT7 (Fig. 1A), Rmo2^HMA+^ bound to PWT7 with nanomolar affinity (Fig. 3B). Similarly, we resurfaced Rwt7^HMA^ to generate Rwt7^HMA+^ (Fig. 3C). Using ITC, we showed that engineered Rwt7^HMA+^ still binds to PWT7, but also gained binding to PBY2 with nanomolar affinity (Fig. 3D). We obtained crystals and determined the structures of Rwt7^HMA+^ co-purified with PBY2 or PWT7, and resolved a tripartite complex containing Rmo2^HMA+^, PBY2, and PWT7 (Fig. 3E–G). The crystal structures of Rmo2^HMA+^ and Rwt7^HMA+^ in complex with their gain-of-binding interacting effectors were consistent with the native interactions (fig. S9, S10 table S1). That Rmo2^HMA+^ can bind two effectors raises the potential of engineering TKP immune receptors to respond to multiple effectors simultaneously.

**Fig. 3.**
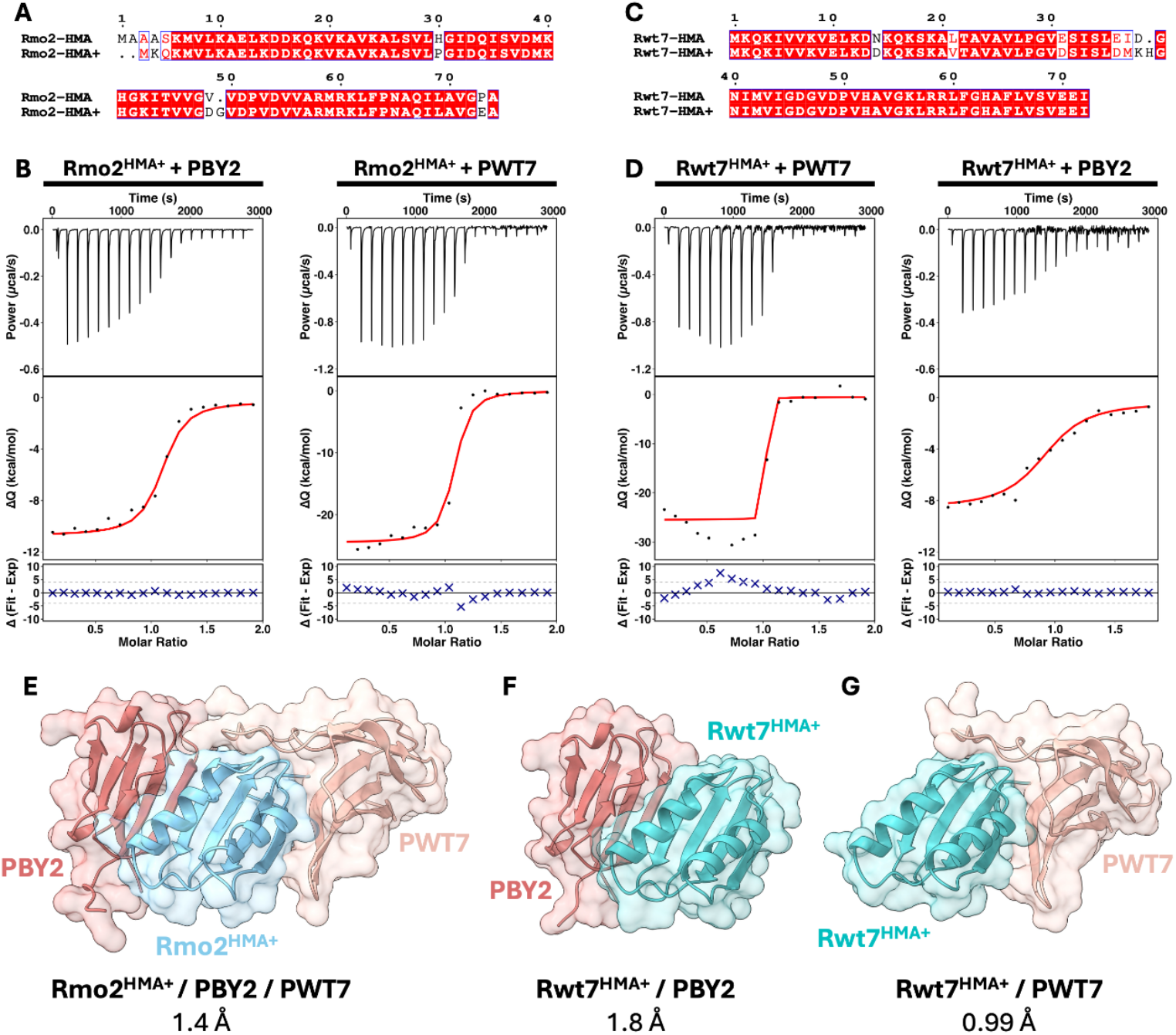
Resurfacing of the integrated HMA domains from TKPs confers binding to non-cognate effectors *in vitro*. (**A**) Amino-acid sequence alignment of Rmo2^HMA^ and the resurfaced Rmo2^HMA+^. (**B**) Binding of Rmo2^HMA+^ to PBY2 or PWT7 by ITC. (**C**) Amino-acid sequence alignment of Rwt7^HMA^ and the resurfaced Rwt7^HMA+^. (**D**) ITC analysis of Rwt7^HMA+^ with PWT7 or PBY2. (**E–G**) Crystal structures of **(E)** the Rmo2^HMA+^/PBY2/PWT7 tripartite complex (PDB: 9TFQ), (**F**) the Rwt7^HMA+^/PBY2 (PDB: 9TFS) co-structure, and (**G**) the Rwt7^HMA+^/PWT7 (PDB: 9TFT) co-structure.

We then tested whether gain of effector binding to Rmo2^HMA+^ or Rwt7^HMA+^ *in vitro* expanded the binding and response of full-length receptors in plant cells. We therefore replaced Rmo^HMA^ with Rmo2^HMA+^ in Rmo2 to generate full-length engineered Rmo2+. Using the split GAL4 RUBY assay in *N. benthamiana*, overall betalain production was comparable when *Rmo2-GAL4* or *Rmo2+-GAL4* was co-expressed with *PBY2-VP16*, suggesting that Rmo2+ still binds to PBY2 *in planta* (Fig. 4, A and B). Importantly, we observed strong betalain production when *Rmo2+-GAL4* was co-expressed with *PWT7-VP16*, in contrast to little betalain accumulation when *Rmo2-GAL4* was co-expressed with *PWT7-VP16*. This finding suggests that Rmo2+ gained binding to PWT7 *in planta*. As Rwt7 is not suitable for the split GAL4 RUBY assay, we used chimeric Rmo2^Rwt7-HMA^ as backbone to replace Rwt7^HMA^ with Rwt7^HMA+^, generating the engineered receptor Rmo2^Rwt7-HMA^+. Overall betalain production in the split GAL4 RUBY assay with *Rmo2*^*Rwt7-HMA*^*+-GAL4* co-expressed with *PWT7-VP16* was comparable to that with *Rmo2*^*Rwt7-HMA*^, suggesting that Rmo2^Rwt7-HMA^+ retains binding to PWT7. Co-expressing *Rmo2*^*Rwt7-HMA*^*+-GAL4* with *PBY2-VP16* also resulted in significant levels of betalain, in contrast to *Rmo2*^*Rwt7-HMA*^*-GAL4* co-expressed with *PBY2-VP16* (Fig. 4, C and D). This result suggests that Rmo2^Rwt7-HMA^+ gained binding to PBY2 *in planta*. When either engineered receptor construct (*Rmo2+-GAL4* or *Rmo2*^*Rwt7-HMA*^*+-GAL4*) was co-expressed with the effector mutant *PBY2*^*E42R*^*-VP16* or *PWT7*^*N39R*^*-VP16*, we measured significantly lower betalain production, clearly establishing that pigment production is associated with direct effector interactions with the HMA domains of the receptors. Taken together, these data demonstrate that the engineered receptors have gained binding to a non-cognate effector in plant cells.

**Fig. 4.**
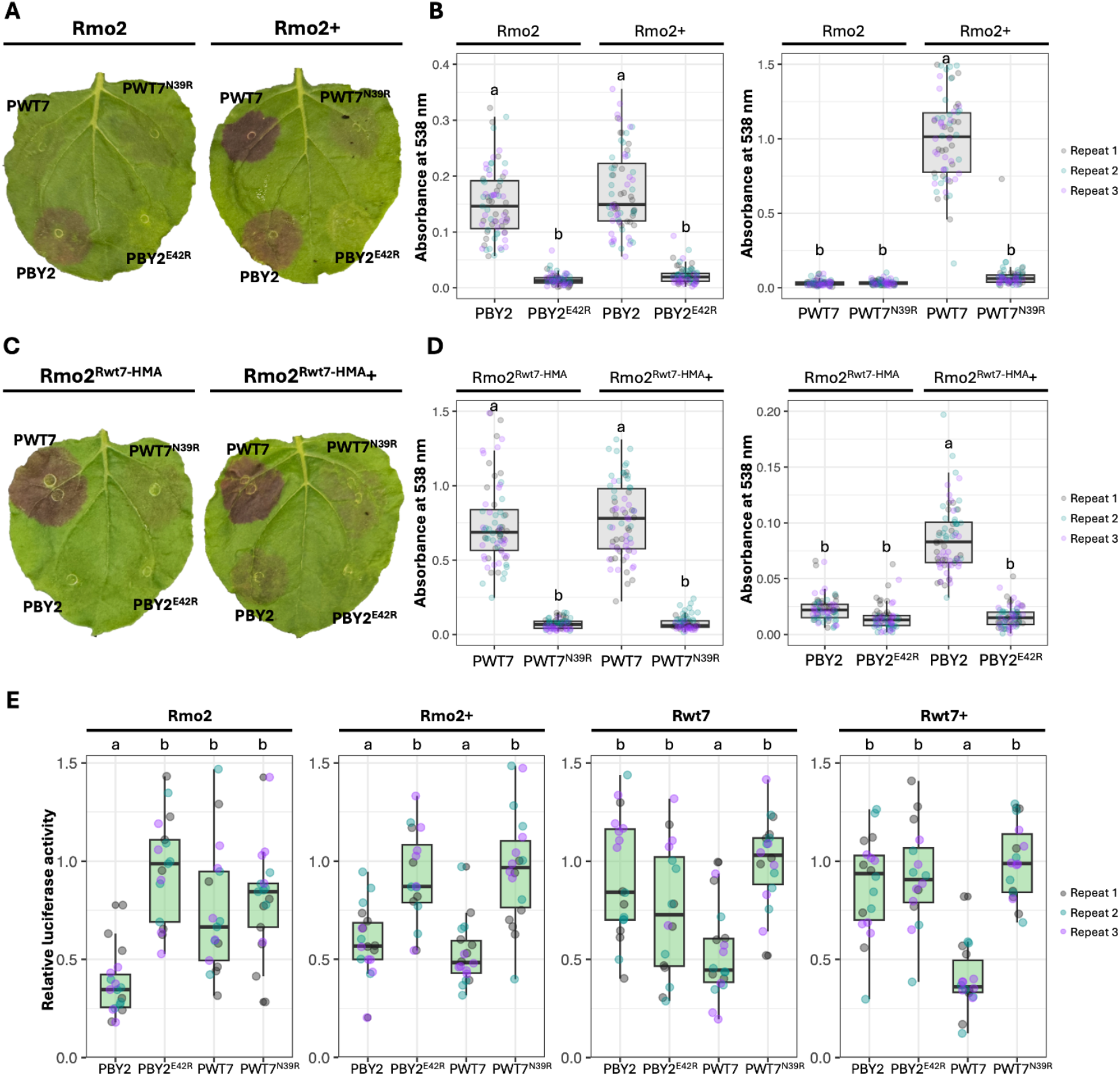
Incorporating the resurfaced HMA domains within TKPs expands their recognition profiles. (**A**) Split GAL4 RUBY assay showing betalain production in *N. benthamiana* leaves following co-expression of *Rmo2-GAL4* or *Rmo2+-GAL4* with *PBY2-VP16, PBY2*^*E42R*^*-VP16, PWT7-VP16*, or *PWT7*^*N39R*^*-VP16*. (**B**) Quantification of betalain production from *N. benthamiana* leaves co-expressing *Rmo2-GAL4* or *Rmo2+-GAL4* with PBY2-VP16, PBY2^E42R^-VP16, PWT7-VP16, or PWT7^N39R^-VP16. (**C**) Split GAL4 RUBY assay showing betalain production in *N. benthamiana* leaves following co-expression of *Rmo2*^*Rwt7-HMA*^*-GAL4* or *Rmo2*^*Rwt7-HMA*^*+-GAL4* with *PBY2-VP16, PBY2*^*E42R*^*-VP16, PWT7-VP16*, or *PWT7*^*N39R*^*-VP16*. (**D**) Betalain quantification of *Rmo2*^*Rwt7-HMA*^*-GAL4* or *Rmo2*^*Rwt7-HMA*^*+-GAL4* co-expressed with *PBY2-VP16, PBY2*^*E42R*^*-VP16, PWT7-VP16*, or *PWT7*^*N39R*^*-VP16*. Leaves were harvested and imaged 4 days after co-infiltration. Leaves were incubated in ethanol until all chlorophyll was cleared. Leaf discs were taken from each infiltration spot after chlorophyll clearing. Betalain was extracted in water and quantified by absorbance at 538 nm. Each experiment consisted of 24 replicates, performed three times independently. Different letters indicate significant differences, as determined by one-way ANOVA and *post-hoc* Tukey’s honestly significant difference tests at *P* < 0.05. (**E**) Relative luciferase activity in wheat protoplasts (from the Kronos cultivar) transfected with constructs encoding Rmo2, Rmo2+, Rwt7, or Rwt7+ in the presence of the effector PBY2 or PWT7, or the effector mutant PBY2^E42R^ or PWT7^N39R^. Luciferase activity is derived from a *proZmUbi:LUC* reporter and was normalized to the values obtained with *PBY2*^*E42R*^ co-expressed with *Rmo2* or *Rmo2*+, and those with *PWT7*^*N39R*^ co-expressed with *Rwt7* or *Rwt7*+. Experiments consisting of six technical replicates were performed three times independently. Different letters indicate significant differences, as determined by one-way ANOVA and *post hoc* Tukey’s honestly significant difference tests at *P* < 0.05.

### Engineered integrated HMA domains expand recognition profiles of TKP receptors in cereal cells

We used a cereal protoplast cell-death assay (*34*) to test whether the gain of effector binding by engineered TKPs was accompanied by expanded recognition and immune responses. In this assay, a reporter construct harboring the firefly luciferase (*LUC*) reporter gene driven by the rice *Ubiquitin* promoter is co-transfected with plasmids encoding relevant receptor/effector combinations into protoplasts prepared from the wheat cultivar ‘Kronos’. Low relative luciferase activity reflects immune activation (cell death). We measured significantly lower relative luciferase activity when constructs encoding intact receptors were co-transfected with constructs encoding the cognate effector (*Rmo2* with *PBY2, Rwt7* with *PWT7*), but not with constructs encoding effector mutants or mismatched effectors (Fig 4E), consistent with the susceptibility/resistance phenotypes during infection assays (Fig 2, C and D). The engineered Rmo2+ and Rwt7+ receptors displayed generally lower luciferase activity compared to wild-type (fig. S11). Nonetheless, protoplasts expressing engineered *Rmo2*+ with either *PBY2* or *PWT7* emitted significantly lower relative luciferase activity than those co-transfected with the non-interacting effector mutants PBY2^E42R^ and PWT7^N39R^ (Fig. 4E), supporting the notion that Rmo2+ gained recognition of PWT7 and retained recognition of PBY2. The engineered Rwt7+ showed significantly lower luciferase activity in the presence of PWT7 but not PBY2, suggesting that although Rwt7^HMA+^ gained binding to PBY2 *in vitro* and *in planta*, this interaction did not translate into a recognition response.

## Discussion

Here, we demonstrate that the integrated HMA domains within the barley TKP Rmo2 and the wheat TKP Rwt7 act as bait domains to directly bind effectors from the blast pathogen and are necessary and sufficient for defining specificity. Structural analyses of the integrated HMA domain/effector complexes revealed interfaces between these proteins that can be altered by mutation, either to prevent effector binding and disease resistance or to gain novel immune signaling profiles.

Although effector-binding integrated domains are an established feature of NLR immune receptors, their presence in TKPs has only recently emerged, leaving their roles underexplored. Extensive genome mining across the plant kingdom indicated that over 56% of TKPs harbor integrated domains (*30*). Importantly, mutations within the integrated domains of TKPs can lead to loss of disease resistance (*19, 27*), demonstrating their importance for protein function. Of the cloned TKPs with integrated HMA domains, the HMA sequences are more divergent compared to their more conserved kinase domains (fig. S1). This observation suggests that these integrated HMA domains are under selective pressure to diversify, potentially to evolve new effector-binding capabilities. Allelic variation is apparent, for example, within *Rpg1* and *Rmo2* across barley cultivars, as alleles of *Rpg1* (*12, 35*) and *Rmo2* (*36*) encode proteins with variable HMA domains that may bind to different cereal pathogen effectors.

While Rmo2 was successfully engineered to recognize PWT7 in a cereal protoplast assay, the engineered Rwt7 failed to recognize PBY2, despite PBY2 interacting with this engineered HMA domain *in vitro* (Fig 3D, Fig 4E). A similar phenomenon was observed with engineered RGA5, which did not lead to resistance in transgenic rice against *M. oryzae* expressing *Avr-PikD*, despite *in vitro* binding of the engineered RGA5 HMA domain to Avr-PikD (*37*). It was observed that the binding affinity of Rwt7^HMA+^ with PBY2 was lower than that of Rwt7^HMA^ with PWT7 and Rmo2^HMA^ with PBY2 (Fig 2, B and D, Fig 3D). Rwt7 may have a higher binding affinity requirement for activation than Rmo2 and the lower binding affinity of PBY2 may be insufficient to activate Rwt7+-mediated immune responses. Another possibility is that PBY2 binds to Rwt7+ in a way that cannot activate the receptor for downstream immune signaling. Elucidating the mechanism of immunity mediated by the HMA domains of TKPs will greatly improve our ability to precisely engineer these receptors in the future.

Many characterized intracellular immune receptors require protein partners to convert pathogen perception into responses. Coiled-coil-type NLRs (CNLs) can be singletons (detect effectors and directly promote cell-death), but many “sensor” CNLs (that perceive effectors) signal through “helper” CNLs encoded by paired genes or as part of genetic networks (*5*). Notably, specific TKPs appear to also require an NLR for immune signaling following effector perception. Sr62^TK^, WTK3, and Rwt4 require the same partner NLR, highlighting how NLRs can act as signaling hubs for multiple TKPs (*25, 26*). Whether Rmo2 and Rwt7 require an NLR to execute immune signaling after effector perception is currently unknown. In Rpg1, the catalytic lysine residues in the pseudokinase (K152) and kinase (K461) domains are both required for disease resistance (*38*), and one susceptible *Rpg1* allele (WBDC323) carries mutations within the pseudokinase domain (I269F, S319RF), suggesting that the pseudokinase domain is important for downstream signaling (*35*), as observed for Sr62^TK^ and Rwt4 (*25, 27*).

Plant immune receptor engineering is an emerging tool to help curb crop losses to plant disease in the future by expanding the recognition profiles of receptors and the generation of new-to-nature disease resistance (*39-41*). Engineering the integrated domains of NLRs for effector recognition has until now been a primary focus and has included domain swapping, structure-informed mutagenesis, directed evolution, and the incorporation of foreign proteins such as nanobodies (*40, 42-46*). In this study, we demonstrate that this approach can be extended to another class of receptors, the TKPs. The stage is now set for studies to explore engineering of the many recently discovered TKPs in diverse cereal crops to determine their potential for developing new-to-nature disease resistance profiles in agriculture.

## Acknowledgements

The authors thank all members of the Banfield Laboratory for discussions. The authors also thank Abbas Maqbool (JIC Biophysical Analysis), David Lawson, Sandra Eltschkner, and Julia Mundy (JIC Structural Biology), Mark Youles and Liam Egan (TSL SynBio) for their expert help and advice in this work, and JIC Horticultural Services for plant growth.

## Funding

UKRI-Biotechnology and Biological Sciences Research Council (UKRI-BBSRC, UK, grants BB/X010996/1 and UKRI1913); John Innes Foundation; Japan Society for the Promotion of Science (grant 22K20580); Kobe University Strategic International Collaborative Research Grant (Type B Fostering Joint Research).

## Author contributions

Conceptualization: DSY, SA, MJB

Methodology: DSY, RZ, EK, HA, LD, NJW, IG, SA, MJB

Investigation: DSY, RZ, EK, HA, LD, NJW, IG, SA

Funding acquisition: SA, MJB

Project administration: DSY, SA, MJB

Supervision: DSY, RZ, SA, MJB

Writing – original draft: DY, SA, MJB

Writing – review & editing: DSY, RZ, EK, HA, LD, NJW, IG, SA, MJB

## Competing interests

Authors declare that they have no competing interests.

## Data and materials availability

All structures have been deposited in the PDB server with the codes 9TFO, 9TFP, 9TFQ, 9TFR, 9TFS, and 9TFT. All other data are provided in the main text or supplementary materials. Detailed material and methods are provided in the supplementary material section. Plasmid constructs used in this study are available on request, but maybe subject to a material transfer agreement.

## Notes

### Competing Interest Statement

The authors have declared no competing interest.

