## Supplemental Information for "Engineering plant tandem kinase immune receptors expands effector recognition profiles"

**The PDF file includes:**

Materials and Methods
Figs. S1 to S11
Tables S1 to S2

### Materials and methods

#### Molecular cloning

All plasmids were generated using the Goldengate system using resources from TSL Synbio (*47*) or Infusion cloning (Takara Bio UK LTD, UK). A list of plasmids constructed in this study and domain boundaries used can be found in table S2.

#### Production and purification of recombinant proteins

Sanger sequencing–verified constructs were chemically transformed into SHuffle T7 Express C3029 cells (NEB) using the protocol provided by the manufacturer and the transformants were grown on LB agar plates containing 100 µg/mL carbenicillin at 37°C for 16 h. For co-expression, plasmids encoding a 6xHis-tagged effector and untagged HMA domain were simultaneously transformed into SHuffle cells and grown on LB agar plates containing 100 µg/mL carbenicillin and 50 µg/mL kanamycin. Colonies were used to inoculate liquid LB containing the required antibiotics; the resulting cultures were grown overnight at 30°C with shaking. Small-scale overnight cultures were used to inoculate 1 L of liquid LB in a 2-L baffled flask to which the required antibiotics were added, together with 2 mM $MgCl_2$ and 200 µL of Antifoam 204 (Sigma-Aldrich Inc., Missouri, USA). These large-scale cultures were incubated at 30°C with shaking at 220 rpm. Cultures were induced to produce the recombinant protein with the addition of isopropyl β-D-1-thiogalactopyranoside (IPTG) to a final concentration of 1 mM once an optical density at 600 nm ($OD_{600}$) of 0.6 was reached, followed by incubation at 18°C with shaking at 220 rpm for 16 h. Cells were harvested by centrifugation at 5000 g for 10 min at 4°C.

Pellets were resuspended in a buffer containing 50 mM HEPES pH 8.0, 300 mM NaCl, 5% (v/v) glycerol, 20 mM imidazole, and 1 mM phenylmethylsulfonyl fluoride (PMSF). The cell pellets were lysed by sonication using an amplitude of 40% (10 seconds on, 20 seconds off). The lysed cells were centrifuged at 30000 g for 30 min at 4°C to clarify the lysate. The protein of interest was purified by immobilized metal affinity chromatography (IMAC) using a 5-mL HisTrap FF nickel column (Cytiva, Massachusetts, USA). The column was washed using a buffer consisting of 50 mM HEPES pH 8.0, 300 mM NaCl, 5% (v/v) glycerol, and 20 mM imidazole prior to elution using an isocratic elution with a buffer composed of 50 mM HEPES pH 8.0, 300 mM NaCl, and 250 mM imidazole. Eluted fractions were analyzed by SDS-PAGE and fractions containing the protein of interest were desalted using a 53-mL SepFast 26/10 1000-6000 Da Desalting Column (BioToolomics Ltd, UK) into 20 mM HEPES pH 7.5, 150 mM NaCl. Fusion proteins were incubated with 3C protease overnight at 4°C; cleavage of the solubility tag was confirmed by SDS-PAGE. The cleaved protein of interest was separated from the N-terminal fusion tag, any uncleaved protein, and 6xHis tagged 3C protease using IMAC and subsequently purified further by size-exclusion chromatography (SEC) using a HiLoad 26/600 Superdex 75 pg column (Cytiva) equilibrated with a buffer consisting of 20 mM HEPES pH 7.5 and 150 mM NaCl. Proteins were concentrated using a 3-kDa or 10-kDa molecular weight cut-off Amicon centrifugal concentrator (Millipore Sigma, Massachusetts, USA), frozen in liquid nitrogen, and stored at −70°C until use.

#### Crystallization and crystal optimization

Initial screening to determine the crystallization conditions for co-purified HMA/effector complexes were performed at a concentration of 15.9 mg/mL for $Rmo2^{HMA}$/PBY2, 16.1 mg/mL for $Rwt7^{HMA}$/PWT7, 14.8 mg/mL for $Rmo2^{HMA+}$/PBY2, 20 mg/mL for $Rmo2^{HMA+}$/PBY2/PWT7,

14.6 mg/mL for Rwt7$^{HMA+}$/PBY2, and 16.7 mg/mL for Rwt7$^{HMA+}$/PWT7, in 96-well MRC 2 plates (Hampton Research, California, USA) with incubation at 18°C using the sitting-drop vapor-diffusion method and commercially available sparse matrix screens. Then, 300 nL of each protein solution and 300 nL of reservoir solution were mixed on a sitting-drop well using an Oryx8 robot (Douglas Instruments, UK) and were monitored and imaged using the Rock Imager system (Formulatrix, Massachusetts, USA) over the course of one month. If required, the original crystal hits were optimized in 96-well MRC 2 sitting-drop plates using an Oryx8 robot (Douglas Instruments)

From initial screening, crystals with the best morphology for Rmo2$^{HMA}$/PBY2 were obtained in 2.4 M ammonium sulfate, 0.1 M sodium acetate, pH 4.0 (C3, KISS, JIC in-house screen). Crystal optimization was carried out and the final condition was in 1.6 M ammonium sulfate, 0.1 M sodium acetate, pH 4.2. For both Rwt7$^{HMA}$/PWT7 and Rmo2$^{HMA+}$/PBY2, the best morphology hit was in 0.2 M ammonium sulfate, 0.1 M sodium acetate, pH 4.0, 20% (w/v) polyethylene glycol (PEG) 3350 (E2, KISS, JIC in-house screen). For Rwt7$^{HMA}$/PWT7, the initial hit was optimized, and the final condition was in 0.2 M ammonium sulfate, 0.1 M sodium acetate, pH 4.2, 16% (w/v) PEG 3350. No optimization was required for Rmo2$^{HMA+}$/PBY2. For Rmo2$^{HMA+}$/PBY2/PWT7, crystals were obtained in 0.1 M KBr, 30% (w/v) PEG 2000 MME (G10, JCSG-plus, Molecular Dimensions); the original hit was optimized under the final condition of 0.15 M KBr, 0.1 M MES pH 6.25, 35% (w/v) PEG 2000 MME. The best crystal hit for Rwt7$^{HMA+}$/PBY2 was in 2.4 M ammonium sulfate, 0.1 M MES, pH 6.0 (C11, KISS, JIC in-house screen), while the best hit for Rwt7$^{HMA+}$/PWT7 was in 0.1 M Tris-HCl pH 8.5, 25% (w/v) PEG 6000 (D9, NeXtal PEGs Suite, Molecular Dimensions). No optimization was required for Rwt7$^{HMA+}$/PBY2 or Rwt7$^{HMA+}$/PWT7.

**Data collection and crystal structure determination**

Before X-ray data collection, crystals were transferred into a cryoprotectant solution consisting of 20% (v/v) ethylene glycol, vitrified in liquid nitrogen, and shipped to Diamond Light Source for X-ray data collection. Diffraction data were collected at Diamond Light Source on the i04 beamline under the proposal mx32728. The data were scaled and merged by the Aimless program in the CCP4i2 software package (*48*). All structures were solved by molecular replacement using PHASER (*49*). High-confidence AlphaFold2 models of PBY2, PWT7, Rmo2$^{HMA}$, and Rwt7$^{HMA}$ were generated using the ColabFold server (*31, 32*) and used as search models for molecular replacement. Models were then refined using the REFMAC program in the CCP4i2 package (*50*; model building between refinement rounds was done in COOT (*51*). X-ray data collection and refinement statistics can be found in table S1. Final structures were deposited in the PDB under accession numbers 9TFO, 9TFP, 9TFQ, 9TFR, 9TFS, and 9TFT.

**Analytical size-exclusion chromatography**

Purified recombinant proteins (Rmo2$^{HMA}$, Rwt7$^{HMA}$, Rwt7$^{HMA}$-6His, PBY2, PBY2$^{E42R}$, PWT7, PWT7$^{N39R}$) were incubated on ice for at least 1 h alone or in pairs. The incubated proteins were then separated on a Superdex 75 Increase 5/150 GL (Cytiva) column pre-equilibrated in a buffer composed of 20 mM HEPES, pH 7.5, 150 mM NaCl. Samples across the peaks were analyzed by SDS-PAGE and Coomassie staining of the gels.

**Isothermal titration calorimetry (ITC)**

For ITC experiments, 300 µL of Rmo2$^{HMA}$, Rwt7$^{HMA}$-6His, Rmo2$^{HMA+}$-6His, or Rwt7$^{HMA+}$-6His at a concentration of 20 µM was loaded into the calorimetric cell. Then, 60 µL of PBY2, PBY2$^{E42R}$, PWT7, or PWT7$^{N39R}$ at 200 µM was loaded into the syringe. Each ITC run was conducted at 25 °C in 20 mM HEPES, pH 7.5, 150 mM NaCl, and consisted of a single 0.5-µL syringe injection followed by 18 syringe injections of 2 µL each at 120-second intervals. ITC experiments were performed three times and the data obtained were analyzed using AFFINImeter ITC software (*52*).

**Protoplast assay**

Protoplasts were isolated and transfected according to Knight, Winfield, Goldson, Bargmann and Borrill (*53*). Briefly, leaves from 8–9-day-old wheat seedlings ('Kronos' *Triticum durum* cultivar) grown in the dark were harvested and cut into 0.5–1-mm strips in plasmolysis buffer (750 mM mannitol, 1 mM $CaCl_2$, 15 mM MES-KOH). Chopped leaves were submerged in enzyme solution (1.5% [w/v] Cellulase RS, 0.5% [w/v] Macerozyme R10, 600 mM mannitol, 10 mM MES-KOH, 1 mM $CaCl_2$, 0.1% [w/v] BSA), vacuum-infiltrated, and incubated in the dark for 4 h. Released protoplasts were filtered using a 40-µm filter mesh and centrifuged at 80 g for 3 min with slow acceleration and deceleration settings. The supernatant was replaced with cold W5 buffer (125 mM $CaCl_2$, 154 mM NaCl, 2 mM MES-KOH, 5 mM KCl) and the cell suspension was incubated in the dark for 30 min. Protoplasts were centrifuged at 80 g for 3 min with slow acceleration and deceleration settings and the supernatant was replaced with MMG buffer (600 mM mannitol, 15 mM $MgCl_2$, 4 mM MES-KOH) to a titer of $3.5 \times 10^5$ protoplasts/mL. For transfection, 200 µL of protoplasts were added to tubes containing 10 µg of TKP and effector constructs and 40 µg of the firefly luciferase reporter under the control of the rice *Ubiquitin* promoter. The same volume of PEG solution (40% [w/v] PEG 4000, 80 mM mannitol, 20 mM $Ca(NO_3)_2$) was incubated with the protoplast and DNA mixture for 15 min. The reaction was stopped by adding W5 buffer. Protoplasts were centrifuged at 80 g for 3 min with slow acceleration and deceleration settings and the supernatant was replaced with PIM buffer (600 mM mannitol, 4 mM MES-KOH, 4 mM KCl, 3 mM $CaCl_2$); the resulting cell suspension was incubated in the dark at room temperature for 18–20 h. Luciferase activity measurements were conducted following a protocol adapted from Saur, Bauer, Lu and Schulze-Lefert (*34*) using a Luciferase Assay System (Promega E1500). Luciferase activity was measured in a plate reader. Each protoplast experiment comprised six technical replicates, and each experiment was performed three times. Relative luciferase activity, normalized against reactions containing PBY2$^{E42R}$ or PWT7$^{N39R}$, was plotted and statistical analysis was performed using one-way analysis of variance (ANOVA) and *post-hoc* Tukey's honestly significant difference tests at $P < 0.05$.

**Split GAL4 RUBY assay**

Agrobacterium (*Agrobacterium tumefaciens*) cell cultures (GV3101) each carrying an effector construct with the sequence encoding the herpes virus VP16 activation domain in-frame and downstream of the effector gene, or a receptor construct carrying the sequence encoding the yeast GAL4 DNA-binding domain in-frame and downstream of the receptor gene, both under the control of the cauliflower mosaic virus (CaMV) 35S promoter, were resuspended in infiltration buffer (10 mM MES, pH 5.6, 10 mM $MgCl_2$, and 200 µM acetosyringone). The cell suspensions were mixed with the final $OD_{600}$ values adjusted to 0.5 for both effector and receptor constructs and 0.1 for an Agrobacterium cell suspension harboring the viral silencing suppressor p19. Leaves were harvested and imaged at 4 days after infiltration. For betalain quantification, leaves were incubated in 100% ethanol until all chlorophyll was removed. Two leaf discs (0.8 cm in diameter) from each infiltration spot were taken and incubated in 500 μL $H_2O$ for 6 h at room temperature. Then, 200 μL of the extracted betalain solution was placed in wells within a Greiner F-bottom clear plate and absorbance was measured at 538 nm in a microplate reader (SPECTROstar Nano, BMG LABTECH, Germany). All experiments were performed three times.

**Infection assay**

Wheat seeds were pre-germinated on wet filter papers. After 24 h, sprouted seeds were sown in the soil in vermiculite supplied with liquid fertilizer in a seedling case (5.5 × 15 × 10 cm) and grown at 22°C in a controlled-environment room under a 12-h light/12-h dark photoperiod provided by fluorescent light bulbs for eight days. Primary leaves from 8-day-old seedlings were fixed onto a hard plastic board with rubber bands just before inoculation. Conidial suspensions ($1 \times 10^5$ conidia per mL) were prepared as described previously (*54*) and sprayed onto fixed primary leaves using an air compressor. The inoculated leaves were incubated in sealed trays under darkness and humid conditions at 22°C for 24 h, then transferred to dry conditions under a 12-h light/12-h dark photoperiod provided by fluorescent light bulbs and incubated at 22°C for an additional 3 days. Four days after inoculation, symptoms were evaluated based on the size and color of lesions (*55*). The size of lesions was rated on a progressive scale from 0 to 5: 0, no visible infection; 1, pinhead spots; 2, small lesions (<1.5 mm); 3, scattered lesions of intermediate size (<3 mm); 4, large typical lesions; and 5, complete blighting of leaf blades. A disease score comprised a number denoting the lesion size and a letter indicating the lesion color: 'B' for brown lesions and 'G' for green lesions. The infection types 0–3 with brown lesions were regarded as resistant (R), while the infection types 3–5 with green lesions were considered susceptible (S).

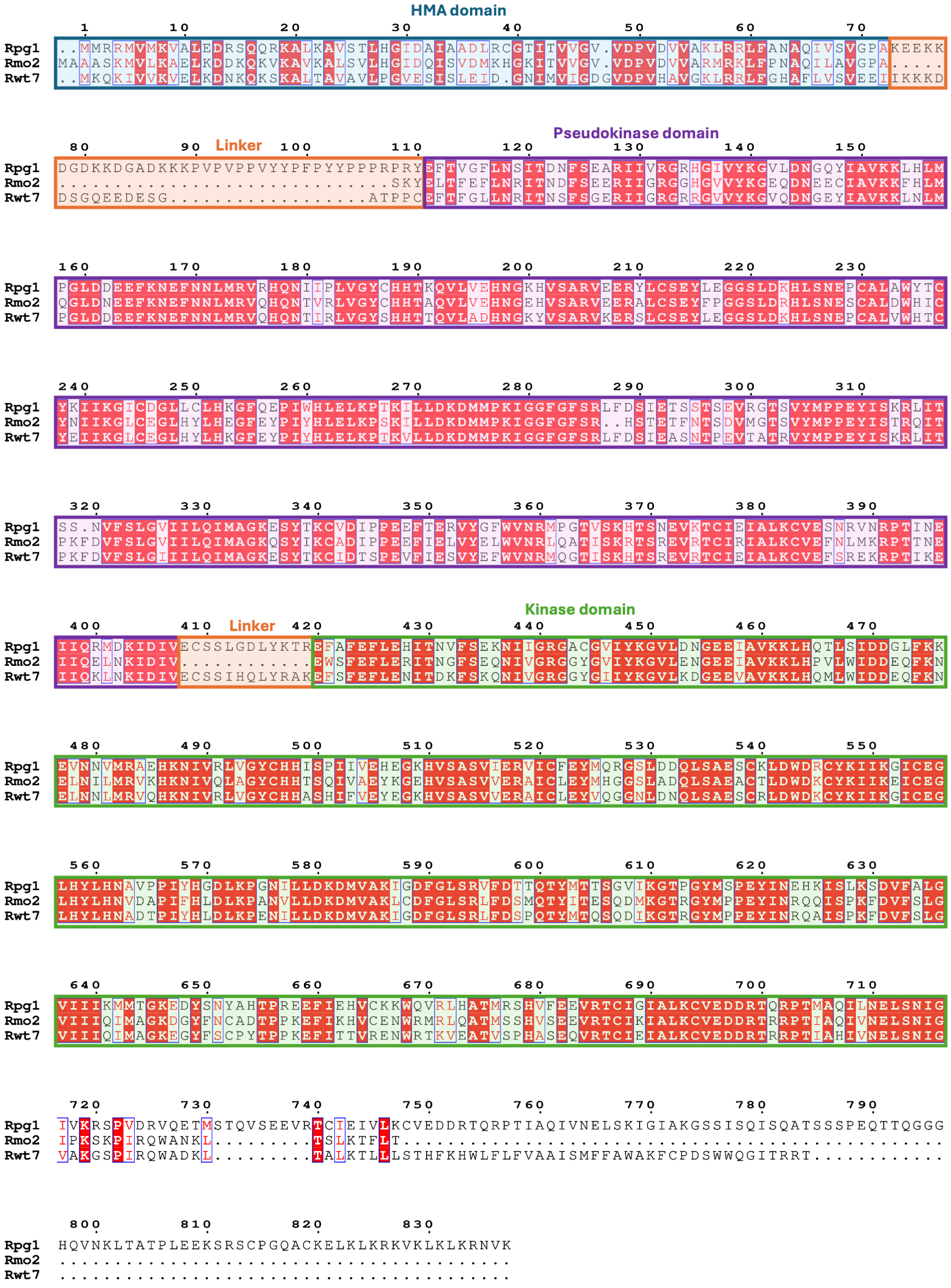


**Fig. S1. Multiple amino-acid sequence alignment of Rmo2, Rwt7 and Rpg1.** The integrated HMA, pseudokinase, and kinase domains are highlighted in blue, purple and green, respectively. Linker regions between the domains are highlighted in orange. The alignment was generated using the ESPript 3.0 server.

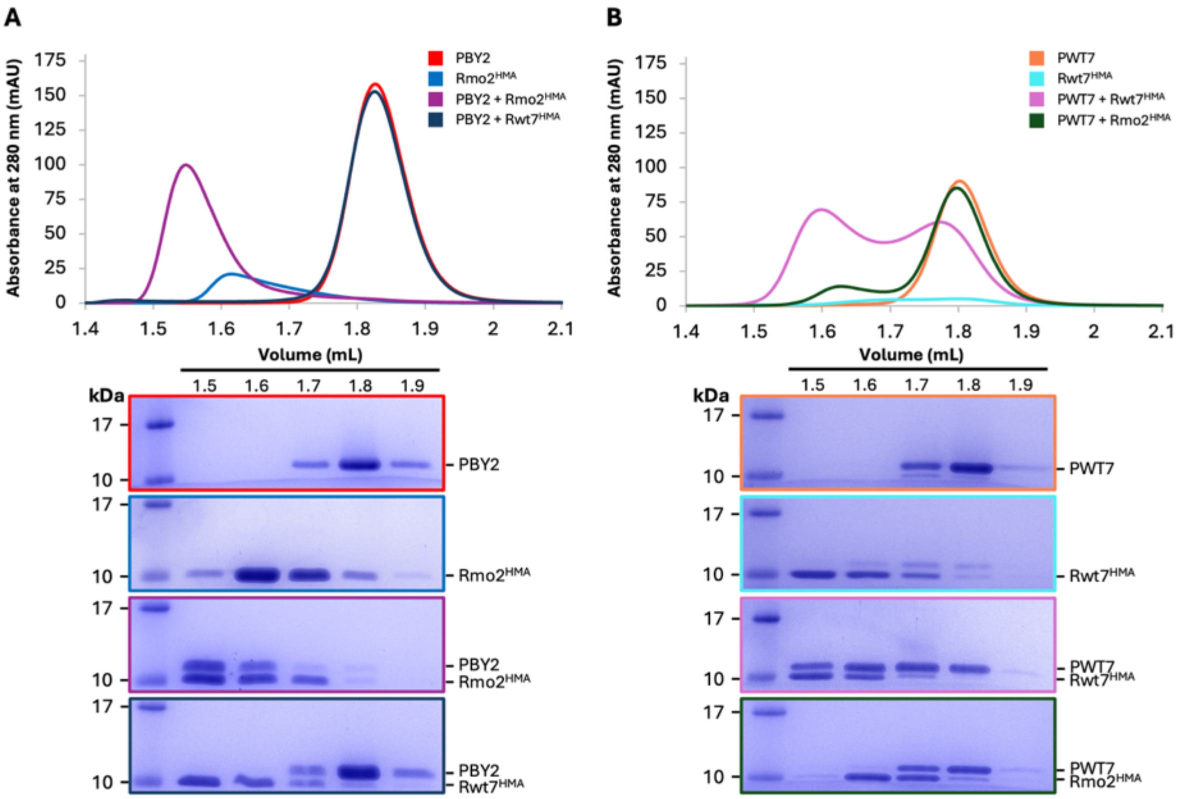


**Fig. S2. Analytical size-exclusion chromatography of integrated HMA domains with their cognate and non-cognate effectors. (A)** Representative size-exclusion chromatograms of PBY2 alone (red), Rmo2$^{HMA}$ alone (blue), PBY2 and Rmo2$^{HMA}$ (purple), PBY2 and Rwt7$^{HMA}$ (navy), and **(B)** PWT7 alone (orange), Rwt7$^{HMA}$ alone (cyan), PWT7 and Rwt7$^{HMA}$ (pink), and PWT7 and Rmo2$^{HMA}$ (dark green). Proteins were incubated on ice for one hour and then separated on a Superdex 75 Increase 5/150 GL column. Coomassie-stained SDS-PAGE gels (bottom panels) depicting samples taken from 100-µL fractions corresponding to the volumes indicated above the gels.

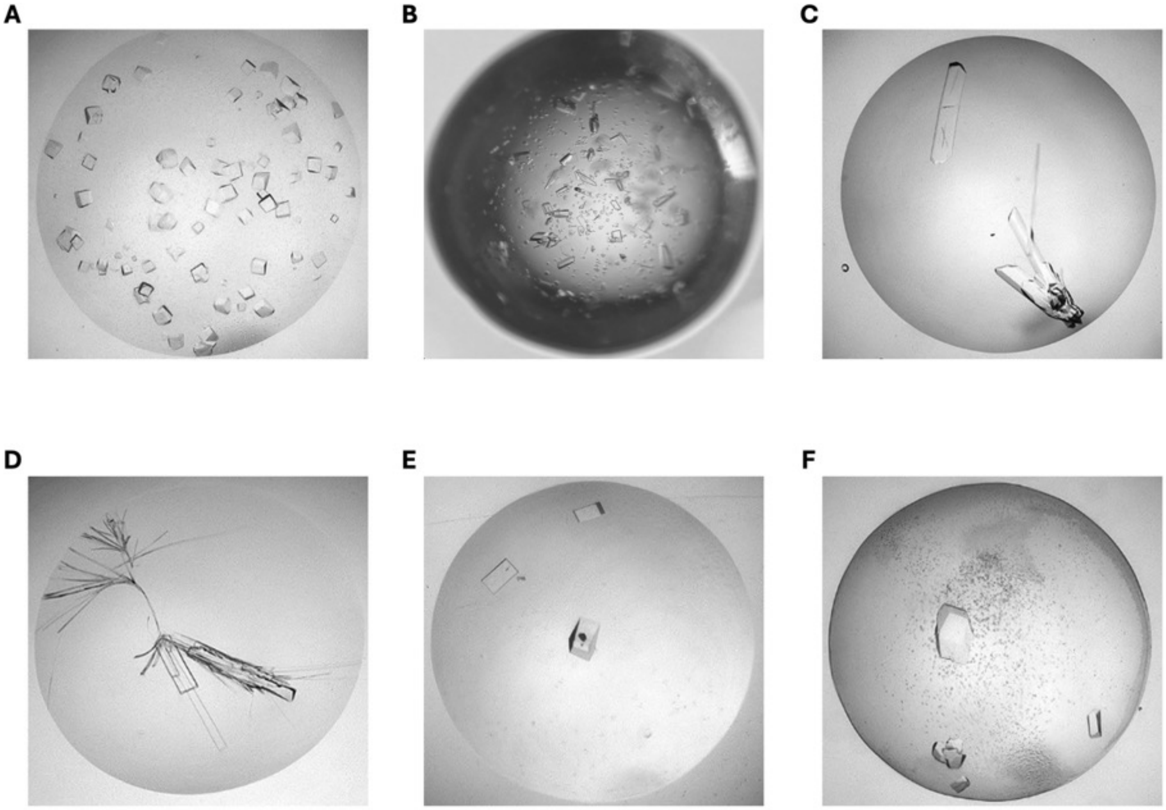


**Fig. S3. Crystallization of HMA/effector complexes.** Optimized protein crystals of **(A)** Rmo2$^{HMA}$/PBY2, **(B)** Rwt7$^{HMA}$/PWT7, **(C)** Rmo2$^{HMA+}$/PBY2, **(D)** Rmo2$^{HMA+}$/PBY2/PWT7, **(E)** Rwt7$^{HMA+}$/PBY2 and **(F)** Rwt7$^{HMA+}$/PWT7.

5

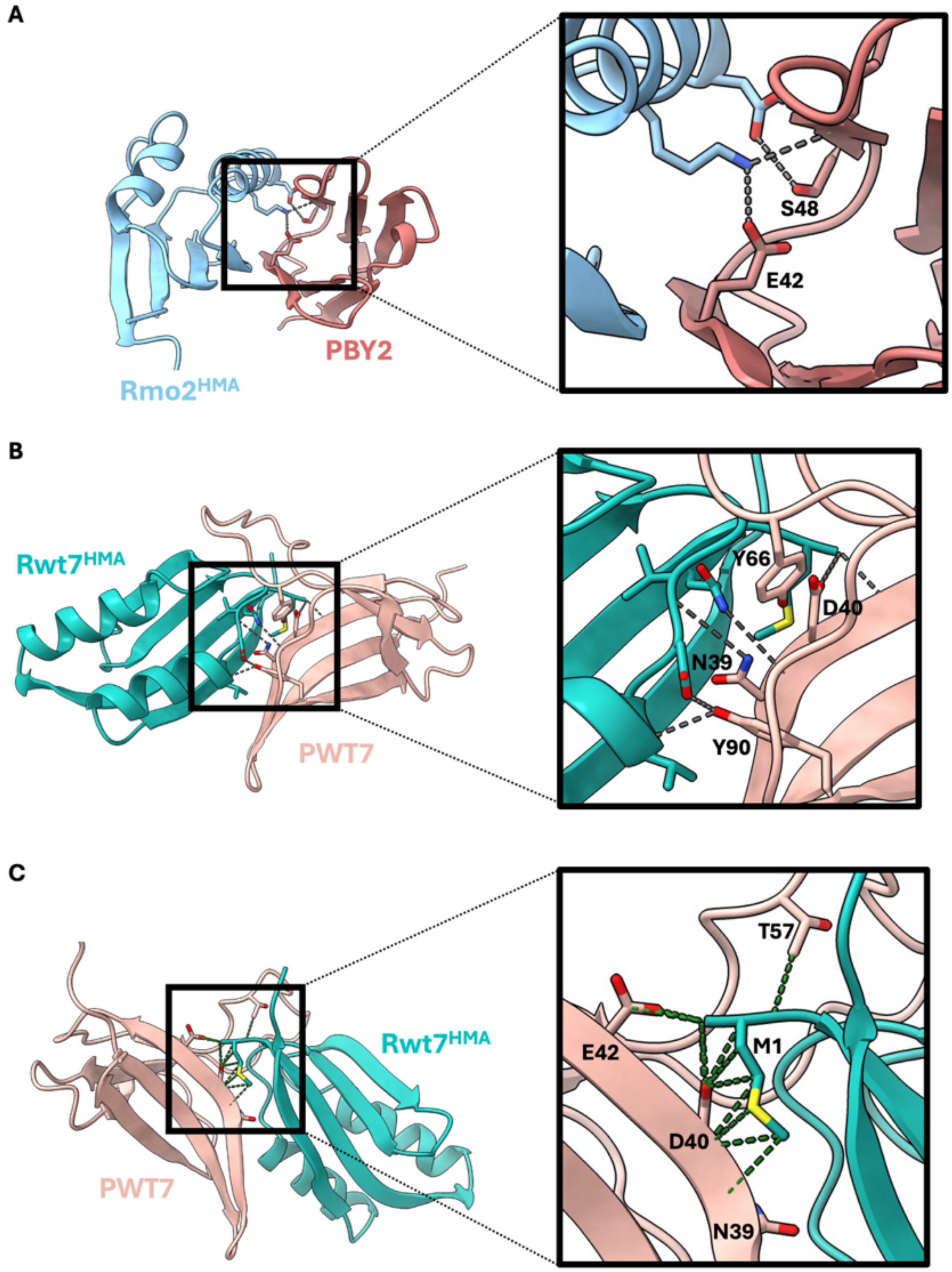


**Fig. S4. Identification of residues within effectors that contribute to their interaction with the integrated HMA domain of Rmo2 or Rwt7. (A)** Hydrogen bonds between Rmo2$^{HMA}$/PBY2 and **(B)** Rwt7$^{HMA}$/PWT7. Interacting residues are shown as sticks. Key residues within the effector proteins are labeled. **(C)** Extensive contacts of the M1 residue from Rwt7$^{HMA}$ with PWT7 residues.

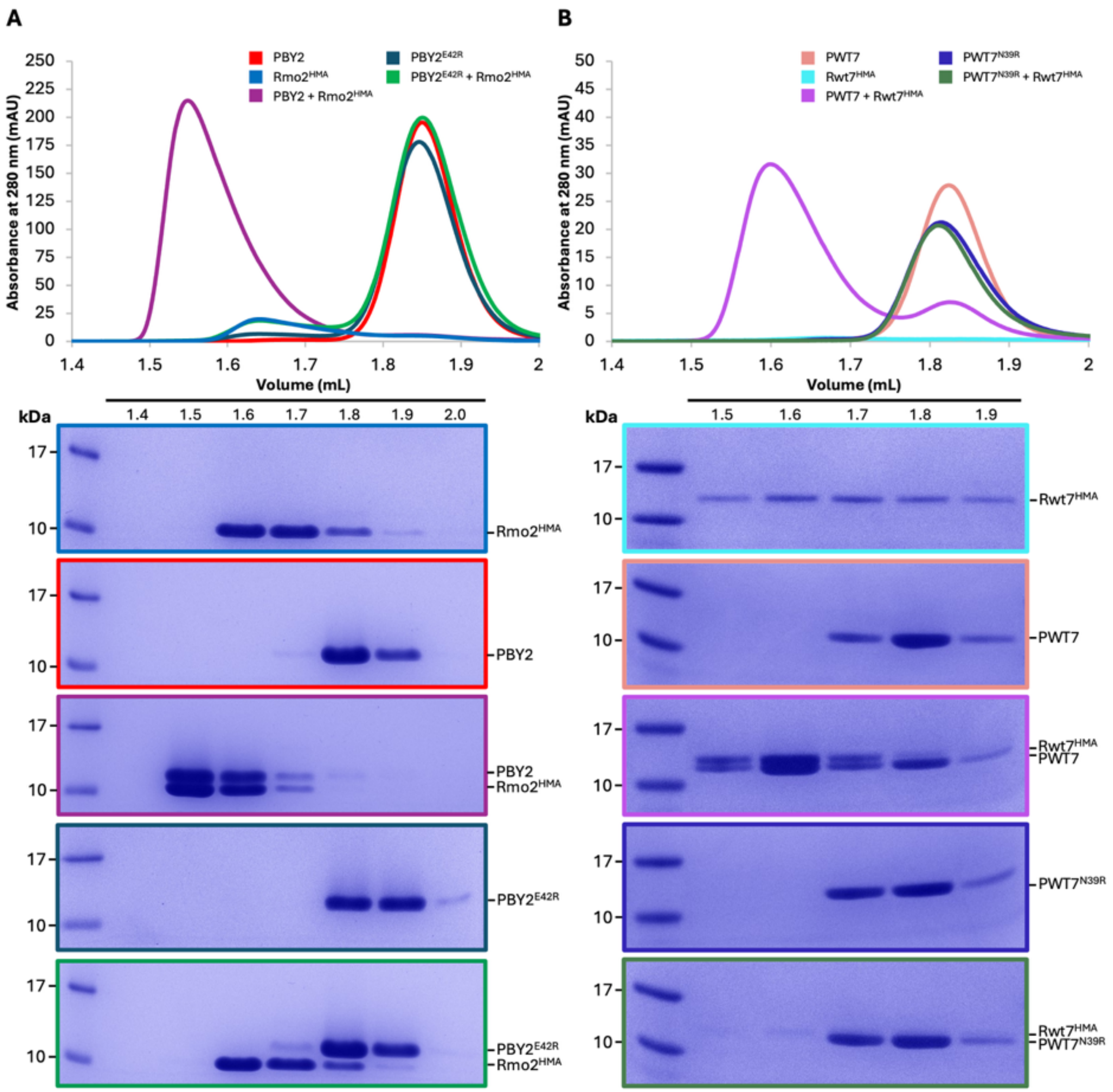


**Fig. S5. Structure-informed effector mutants do not interact with integrated HMA domains by analytical size-exclusion chromatography. (A)** Size-exclusion chromatograms of Rmo2$^{HMA}$ alone (blue), PBY2 alone (red), Rmo2$^{HMA}$ and PBY2 (purple), PBY2$^{E42R}$ alone (navy), Rmo2$^{HMA}$ and PBY2$^{E42R}$ (green), and **(B)** Rwt7$^{HMA}$ alone (cyan), PWT7 alone (orange), Rwt7$^{HMA}$ and PWT7 (pink), PWT7$^{N39R}$ alone (dark blue) and Rwt7$^{HMA}$ and PWT7$^{N39R}$ (dark green). Proteins were incubated on ice for one hour and then separated on a Superdex 75 Increase 5/150 GL column. Coomassie-stained SDS-PAGE gels (bottom panels) depicting samples taken from 100-µL fractions corresponding to the volumes indicated above the gels.

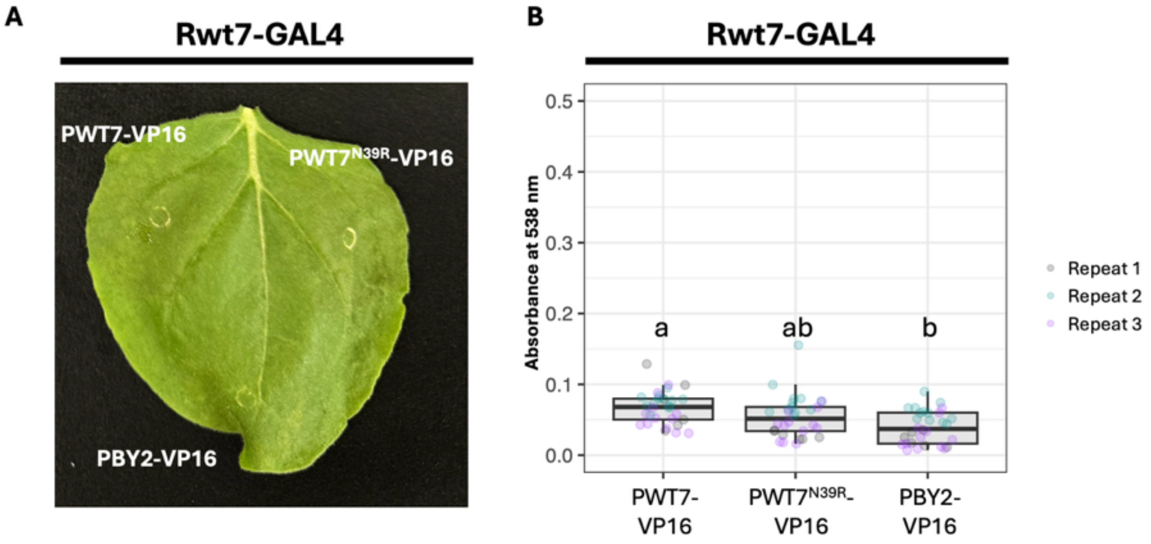


**Fig. S6. Limited betalain is produced with Rwt7-GAL4 using the split GAL4 RUBY assay. (A)** Representative *Nicotiana benthamiana* leaves, showing little betalain production following transient co-expression of *Rwt7-GAL4* (encoding a fusion of Rwt7 with the DNA-binding domain of yeast GAL4 at its C terminus) with *PWT7-VP16*, *PWT7*$^{N39R}$*-VP16*, or *PBY2-VP16* (encoding each effector fused to VP16 at their C termini). All expression cassettes were under the control of the cauliflower mosaic virus (CaMV) 35S promoter. Leaves were harvested 4 days after co-infiltration and chlorophyll was cleared with ethanol. Betalain was extracted from each infiltrated spot and absorbance was measured at 538 nm. Each experiment consisted of 6 or 12 replicates, performed three times independently. Different letters indicate significant differences, as determined by one-way ANOVA and *post-hoc* Tukey's honestly significant difference tests at $P < 0.05$.

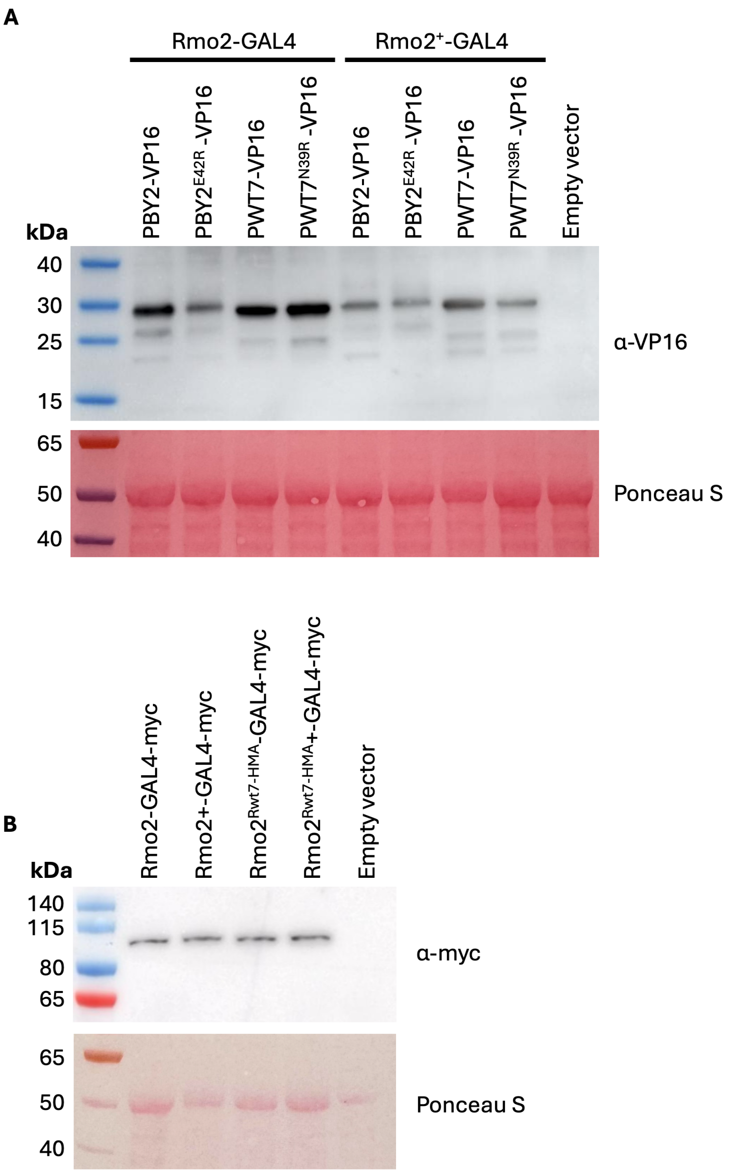


**Fig. S7. Confirmation of protein accumulation from the split GAL4 RUBY constructs in *N. benthamiana*.** Immunoblot analysis of **(A)** PBY2, PBY2$^{E42R}$, PWT7, and PWT7$^{N39R}$ (fused to VP16 at their C termini) in *N. benthamiana* leaves co-infiltrated with the respective encoding constructs and a construct encoding Rmo2 or Rmo2+ (fused to the DNA-binding domain of GAL4 at their C termini), and **(B)** Rmo2, Rmo2+, Rmo2$^{Rwt7\text{-}HMA}$, Rmo2$^{Rwt7\text{-}HMA}$+ (fused to GAL4-myc at their C termini). Total proteins were extracted from *N. benthamiana* leaves 4 days after infiltration and separated by SDS-PAGE. The samples were transferred onto a membrane, probed with anti-VP16 or anti-myc antibodies, and signals were detected with a chemiluminescence imager. The membrane was stained with Ponceau S to show equal sample loading.

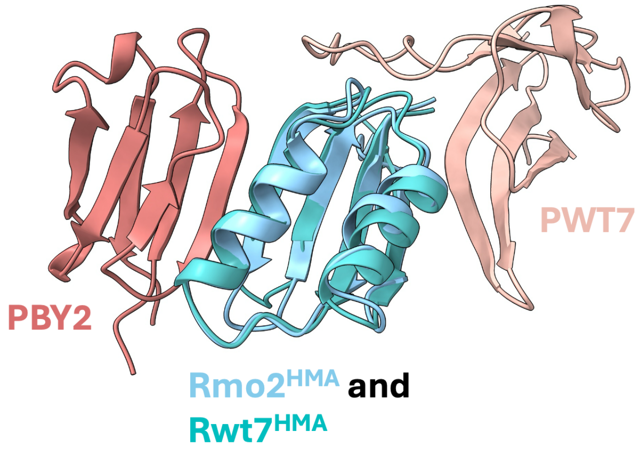


**Fig. S8. Structural alignment of the Rmo2$^{HMA}$/PBY2 and Rwt7$^{HMA}$/PWT7 complexes.** The structures were aligned in ChimeraX.

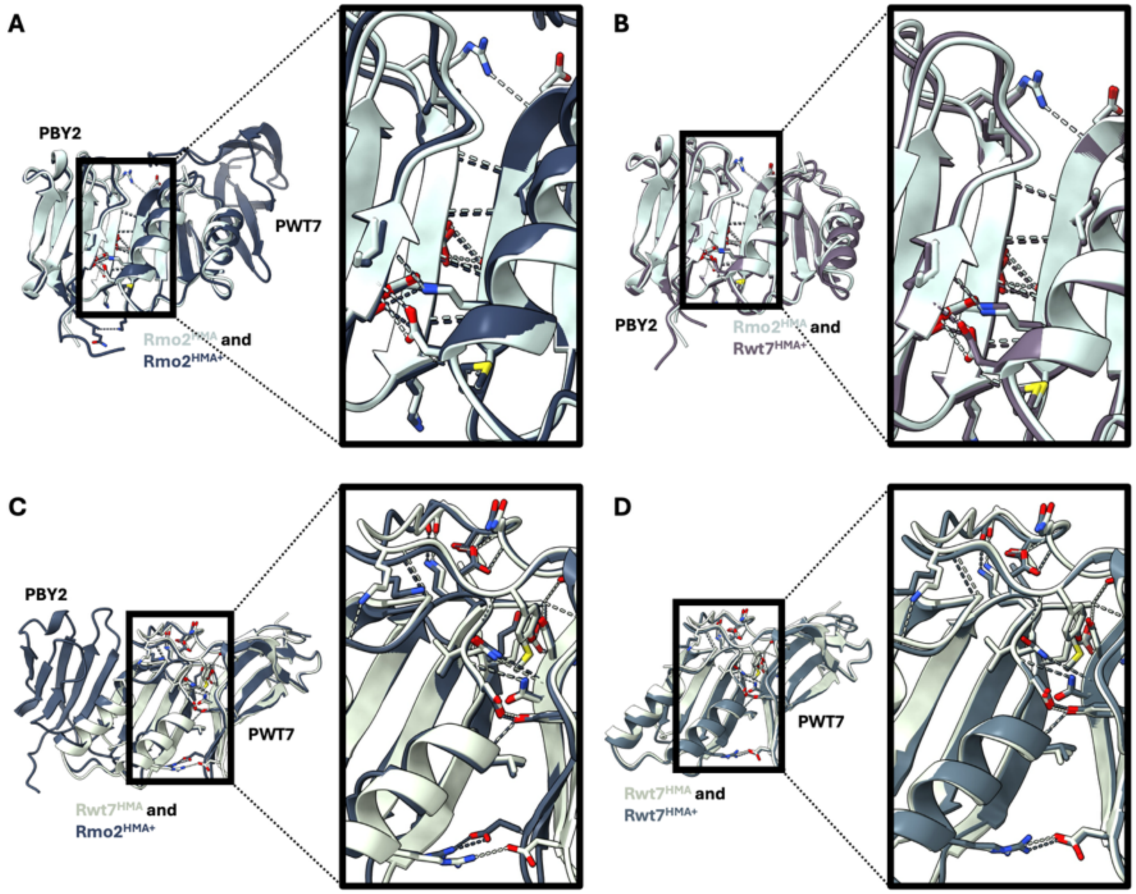


**Fig. S9. Engineered gain-of-binding HMA domains have near identical interaction interfaces.** **(A–D)** Structural alignment of Rmo2$^{HMA}$/PBY2 with **(A)** Rmo2$^{HMA+}$/PBY2/PWT7 or **(B)** Rwt7$^{HMA+}$/PBY2, and of Rwt7$^{HMA}$/PWT7 with **(C)** Rmo2$^{HMA+}$/PBY2/PWT7 or **(D)** Rwt7$^{HMA+}$/PWT7. Wild-type structures are shown in light colors and engineered structures are shown in dark colors. Residues involved in hydrogen bonds are shown as sticks. Hydrogen bonds are shown as dotted lines in light or dark colors from the wild-type or engineered structures, respectively. All structures were aligned in ChimeraX.

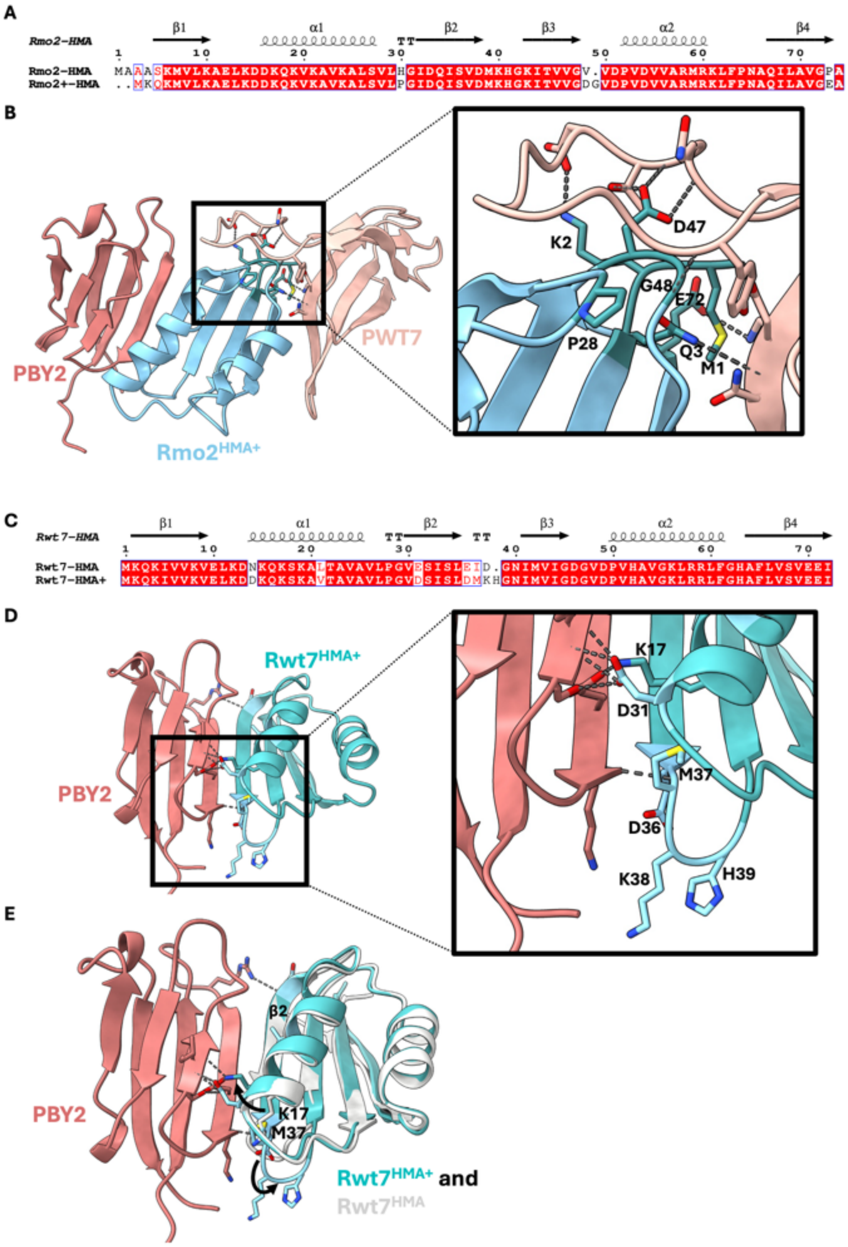


**Fig. S10. All residues from the resurfaced HMAs contribute to gain of binding to non-cognate effectors. (A)** Amino-acid sequence alignment of Rmo2$^{HMA}$ and Rmo2$^{HMA+}$. **(B)** All mutated residues from Rmo2$^{HMA+}$ and corresponding interactions with PWT7. **(C)** Amino-acid sequence alignment of Rwt7$^{HMA}$ and Rwt7$^{HMA+}$. **(D)** Mutated residues from Rwt7$^{HMA+}$ and corresponding interactions with PBY2. **(E)** Resurfacing Rwt7$^{HMA+}$ changes its shape, which facilitates its binding to PBY2. The addition of M37 in Rwt7$^{HMA+}$ displaces K17, which normally forms hydrogen bonds with PBY2. The addition of K38 and H39 straightens out β2 to facilitate extensive main chain hydrogen bonds with PBY2. Mutated residues are labeled and shown as sticks. Hydrogen bonds are shown as dotted lines.

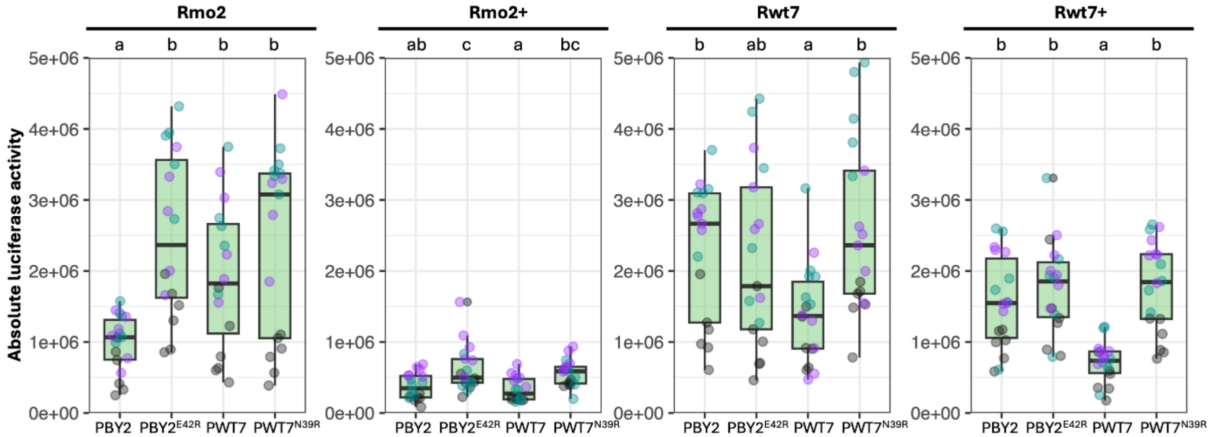


**Fig. S11. Lower overall luciferase activity observed with Rmo2+ and Rwt7+ in wheat protoplasts.** Absolute luciferase activity in wheat protoplasts (from the Kronos cultivar) transfected with constructs encoding Rmo2, Rmo2+, Rwt7, or Rwt7+ in the presence of the effector PBY2 or PWT7, or the effector variant PBY2$^{E42R}$ or PWT7$^{N39R}$. Luciferase activity is derived from a *proZmUbi:LUC* reporter. Experiments consisting of six technical replicates were performed three times independently. Different letters indicate significant differences, as determined by one-way ANOVA and *post-hoc* Tukey's honestly significant difference tests at $P < 0.05$.

**Table S1.** X-ray data collection, structure solution and refinement statistics for PBY2/Rmo2$^{HMA}$, PWT7/Rwt7$^{HMA}$, PBY2/PWT7/Rmo2$^{HMA+}$, PBY2/Rmo2$^{HMA+}$, PBY2/Rwt7$^{HMA+}$, and PWT7/Rwt7$^{HMA+}$.

| | **PBY2/Rmo2$^{HMA}$ (9TFO)** | **PWT7/Rwt7$^{HMA}$ (9TFP)** | **PBY2/PWT7/ Rmo2$^{HMA+}$ (9TFQ)** | **PBY2/Rmo2$^{HMA+}$ (9TFR)** | **PBY2/Rwt7$^{HMA+}$ (9TFS)** | **PWT7/Rwt7$^{HMA+}$ (9TFT)** |
|---|---|---|---|---|---|---|
| Detector | Dectris Eiger2 XE 16M | Dectris Eiger2 XE 16M | Dectris Eiger2 XE 16M | Dectris Eiger2 XE 16M | Dectris Eiger2 XE 16M | Dectris Eiger2 XE 16M |
| Wavelength (Å) | 0.953731 | 0.953738 | 0.953727 | 0.953727 | 0.953709 | 0.750006 |
| Space group | P 31 1 2 | P 2 21 21 | P 1 21 1 | P 31 1 2 | P 43 21 2 | P 1 21 1 |
| Unit cell | 89.792 89.792 89.088 90.000 90.000 120.000 | 39.915 73.609 127.544 90.000 90.000 90.000 | 56.592 30.590 61.448 90.000 93.015 90.000 | 90.075 90.075 88.228 90.000 90.000 120.000 | 105.197 105.197 69.134 90.000 90.000 90.000 | 40.524 66.174 65.747 90.000 104.851 90.000 |
| Average mosaicity (°) [b] | 0.00 | 0.00 | 0.00 | 0.00 | 0.00 | 0.00 |
| Resolution (Å) | 89.09–2.20 (2.27–2.20) | 63.85–1.60 (1.63–1.60) | 61.36–1.40 (1.42–1.40) | 88.23–2.60 (2.72–2.60) | 74.39–1.80 (1.84–1.80) | 66.17–0.99 (1.01–0.99) |
| Total no. of reflections | 438,495 (38,796) | 1,364,358 (67,667) | 566,095 (27,819) | 531,069 (66,239) | 1,680,137 (96,874) | 2,555,845 (125,512) |
| No. of unique reflections | 21,089 (1,838) | 50,590 (2,424) | 41,916 (2,032) | 12,793 (1,545) | 36,570 (2,113) | 186,021 (9,174) |
| Completeness (%) | 100.0 (100.0) | 100.0 (100.0) | 100.0 (100.0) | 100.0 (100.0) | 100.0 (100.0) | 100.0 (100.0) |
| Multiplicity | 20.8 (21.1) | 27.0 (27.9) | 13.5 (13.7) | 41.5 (42.9) | 45.9 (45.8) | 13.7 (13.7) |
| Mean I /s (I) | 26.6 (1.7) | 32.7 (2.0) | 19.8 (1.8) | 24.7 (2.0) | 22.2 (3.9) | 15.2 (1.5) |
| Rmerge | 0.060 (0.931) | 0.052 (1.025) | 0.054 (1.287) | 0.106 (2.412) | 0.145 (1.227) | 0.062 (1.610) |
| Rmeas [c] | 0.063 (0.977) | 0.054 (1.063) | 0.058 (1.386) | 0.108 (2.469) | 0.148 (1.254) | 0.066 (1.740) |
| Rpim [d] | 0.019 (0.295) | 0.014 (0.280) | 0.022 (0.514) | 0.023 (0.526) | 0.030 (0.256) | 0.025 (0.657) |
| $CC_{1/2}$ [b] | 1.000 (0.944) | 1.000 (0.961) | 1.000 (0.872) | 1.000 (0.888) | 1.000 (0.971) | 1.000 (0.817) |
| Matthews coeff. (Å$^3$ Da$^{-1}$) [e] | 2.16 | 2.67 | 2.10 | 3.34 | 3.21 | 2.46 |
| Resolution range (Å) | 77.88–2.20 | 63.85–1.60 | 61.36–1.40 | 78.13–2.60 | 74.50 –1.80 | 63.63–0.99 |
| $R_{work}$ (%) [g] | 0.213 | 0.161 | 0.154 | 0.188 | 0.173 | 0.140 |
| $R_{free}$ (%) [h] | 0.267 | 0.198 | 0.193 | 0.262 | 0.210 | 0.153 |
| No. of non-H atoms | | | | | | |
| Amino acids | 2112 | 2413 | 1737 | 2056 | 2144 | 2503 |
| Ligand | 9 | 64 | 12 | 0 | 89 | 0 |
| Water | 65 | 287 | 194 | 26 | 305 | 477 |
| Average B-factor (Å$^2$) | 75.581 | 37.751 | 26.850 | 93.148 | 26.834 | 14.631 |
| RMSD from ideal geometry | | | | | | |
| Bond lengths (Å) | 0.0071 | 0.0103 | 0.0107 | 0.0130 | 0.0112 | 0.0125 |
| Bond angles (°) | 1.796 | 1.776 | 1.929 | 2.772 | 1.893 | 1.948 |
| Ramachandran plot, residues in (%)[i] | | | | | | |
| Favored regions | 95.13 | 99.00 | 99.05 | 93.85 | 99.25 | 98.34 |
| Allowed regions | 3.75 | 1.00 | 0.95 | 5.77 | 0.75 | 1.66 |
| Outlier regions | 1.12 | 0.00 | 0.00 | 0.38 | 0.00 | 0.00 |

[a] The values in parentheses are for the highest-resolution shell.
[b] Calculated with AIMLESS.
[c] Rmeas = Σhkl (N(hkl) / [N(hkl) – 1])^(1/2) Σi | Ii(hkl) – (1 / N(hkl)) Σj Ij(hkl) | / Σhkl Σi Ii(hkl), where Ii(hkl) is the intensity of the *i*th measurement of an equivalent reflection with indices hkl.
[d] Rpim = Σhkl (1 / [N(hkl) – 1])^(1/2) Σi | Ii(hkl) – (1 / N(hkl)) Σj Ij(hkl) | / Σhkl Σi Ii(hkl).
[e] Calculated with MATTHEWS_COEF within the CCP4 suite.
[f] Generated by Crank pipeline in the CCP4 suite.
[g] Rwork = Σall | Fobs – Fcalc | / Σall | Fobs |, where Fobs and Fcalc are the observed and calculated structure factor amplitudes.
[h] Rfree is equivalent to Rwork but calculated with reflections (5%) omitted from the refinement process.
[i] Calculated with MolProbity.

### Table S2. Constructs used in this study

| Construct | Domain boundary | Backbone | Purpose |
|---|---|---|---|
| GB1-PBY2 | $PBY2^{21–85}$ | pPGN-C | *E. coli* production, analytical SEC, ITC, structures |
| GB1-PWT7 | $PWT7^{22–98}$ | pPGN-C | *E. coli* production, analytical SEC, ITC, structures |
| GB1-$Rmo2^{HMA}$ | $Rmo2^{1–77}$ | pPGN-C | *E. coli* production, analytical SEC, ITC |
| GB1-$Rwt7^{HMA}$ | $Rwt7^{1–77}$ | pPGN-C | *E. coli* production, analytical SEC |
| $Rmo2^{HMA}$ | $Rmo2^{1–77}$ | pPGC-K | *E. coli* production, structures |
| $Rwt7^{HMA}$ | $Rwt7^{1–77}$ | pPGC-K | *E. coli* production, structures |
| GB1-$PBY2^{E42R}$ | $PBY2^{21–85}$ | pPGN-C | *E. coli* production, analytical SEC, ITC |
| GB1-$PWT7^{N39R}$ | $PWT7^{22–98}$ | pPGN-C | *E. coli* production, analytical SEC, ITC |
| Rwt7-6His | $Rwt7^{1–77}$ | pPGC-C | *E. coli* production, ITC |
| $Rmo2^{HMA+}$-6His | $Rmo2^{1–77}$ | pPGC-C | *E. coli* production, ITC |
| $Rwt7^{HMA+}$-6His | $Rwt7^{1–77}$ | pPGC-C | *E. coli* production, ITC |
| $Rmo2^{HMA+}$ | $Rmo2^{1–77}$ | pPGC-K | *E. coli* production, structures |
| $Rwt7^{HMA+}$ | $Rwt7^{1–77}$ | pPGC-K | *E. coli* production, structures |
| 35S:Rmo2-GAL4:T35S | $Rmo2^{1–692}$ | pICSL22027 | RUBY assay |
| 35S:Rwt7-GAL4:T35S | $Rwt7^{1–758}$ | pICSL22027 | RUBY assay |
| 35S:$Rmo2^{Rwt7\text{-}HMA}$-GAL4:T35S | $Rwt7^{1–72}$; $Rmo2^{75–692}$ | pICSL22027 | RUBY assay |
| 35S:Rmo2+-GAL4:T35S | $Rmo2^{1–692}$ | pICSL22027 | RUBY assay |
| 35S:$Rmo2^{Rwt7+\text{-}HMA}$-GAL4:T35S | $Rwt7^{1–72}$; $Rmo2^{75–692}$ | pICSL22027 | RUBY assay |
| 35S:PBY2-VP16:T35S | $PBY2^{21–85}$ | pICSL22026 | RUBY assay |
| 35S:$PBY2^{E42R}$-VP16:T35S | $PBY2^{21–85}$ | pICSL22026 | RUBY assay |
| 35S:PWT7-VP16:T35S | $PWT7^{22–98}$ | pICSL22026 | RUBY assay |
| 35S:$PWT7^{N39R}$-VP16:T35S | $PWT7^{22–98}$ | pICSL22026 | RUBY assay |
| pZH2Bik-PBY2 | $PBY2^{21–85}$ | pZH2Bik | Protoplast assay |
| pZH2Bik-$PBY2^{E42R}$ | $PBY2^{21–85}$ | pZH2Bik | Protoplast assay |
| pZH2Bik-PWT7 | $PWT7^{22–98}$ | pZH2Bik | Protoplast assay |
| pZH2Bik-$PWT7^{N39R}$ | $PWT7^{22–98}$ | pZH2Bik | Protoplast assay |
| pZH2Bik-Rmo2 | $Rmo2^{1–692}$ | pZH2Bik | Protoplast assay |
| pZH2Bik-Rmo2+ | $Rmo2^{1–692}$ | pZH2Bik | Protoplast assay |
| pZH2Bik-Rwt7 | $Rwt7^{1–758}$ | pZH2Bik | Protoplast assay |
| pZH2Bik-Rwt7+ | $Rwt7^{1–758}$ | pZH2Bik | Protoplast assay |
